# Comparative analysis of drug-salt-polymer interactions by experiment and molecular simulation improves biopharmaceutical performance

**DOI:** 10.1101/2022.08.11.503409

**Authors:** Sumit Mukesh, Goutam Mukherjee, Ridhima Singh, Nathan Steenbuck, Carolina Demidova, Prachi Joshi, Abhay T. Sangamwar, Rebecca C. Wade

## Abstract

The propensity of poorly water-soluble drugs to aggregate at supersaturation impedes their bioavailability. The emergence of supersaturated amorphous drug-salt-polymer systems provides a new approach to this problem. However, the effects of polymers on drug-drug interactions in aqueous phase are largely unexplored and it is unclear how to choose an optimal salt-polymer combination for a particular drug. We describe a comparative experimental and computational characterization of amorphous solid dispersions containing the drug celecoxib, and PVP-VA or HPMCAS polymers with or without Na^+^/K^+^ salts. Classical models for drug-polymer interactions fail to identify the best drug-salt-polymer combination. In contrast, more stable drug-polymer interaction energies computed from molecular dynamics simulations correlate with prolonged stability of supersaturated amorphous drug-salt-polymer systems, along with better dissolution and pharmacokinetic profiles. The celecoxib-salt-PVP-VA formulations exhibit excellent biopharmaceutical performance, offering the prospect of less frequent administration and lower doses of this widely used anti-inflammatory, thereby increasing cost-effectiveness, and reducing side-effects.

## INTRODUCTION

Many pharmaceutical drugs are poorly water-soluble, and this property hinders their ability to reach the systemic circulation in the required concentration for optimal therapeutic effect. To overcome this problem, such drugs can be formulated in a high-energy amorphous solid state. This allows attainment of supersaturation exceeding the equilibrium solubility of the drug and thereby increases the flux across biological membranes^[1–3]^. However, the supersaturated state of drug molecules is metastable and therefore the drug molecules tend to aggregate and recrystallize, adversely affecting the biopharmaceutical properties of the drug^[4–6]^. In a typical pipeline to select excipients, such as polymers^[7–9]^, lipids^[10–12]^ or surfactants^[13–15]^, to attain supersaturation of a poorly water-soluble drug (PWSD) and thereby enhance bioavailability, various supersaturated drug delivery systems are explored. A better understanding of the determinants of drug-drug and drug-excipient interactions and their relation to drug bioavailability would allow a more rational choice of drug-excipient combinations. Towards this goal, we here describe a comparative computational and experimental investigation of polymeric excipients for celecoxib (CEL) (Fig. 1A), a selective COX-2 inhibitor that is a biopharmaceutical classification system (BCS) class II PWSD. CEL is widely used for treating inflammatory diseases, including rheumatoid arthritis, osteoarthritis and ankylosing spondylitis,^[16–18]^ and hence a formulation with better solubility and bioavailability would be of enormous therapeutic benefit.

**Fig. 1.**
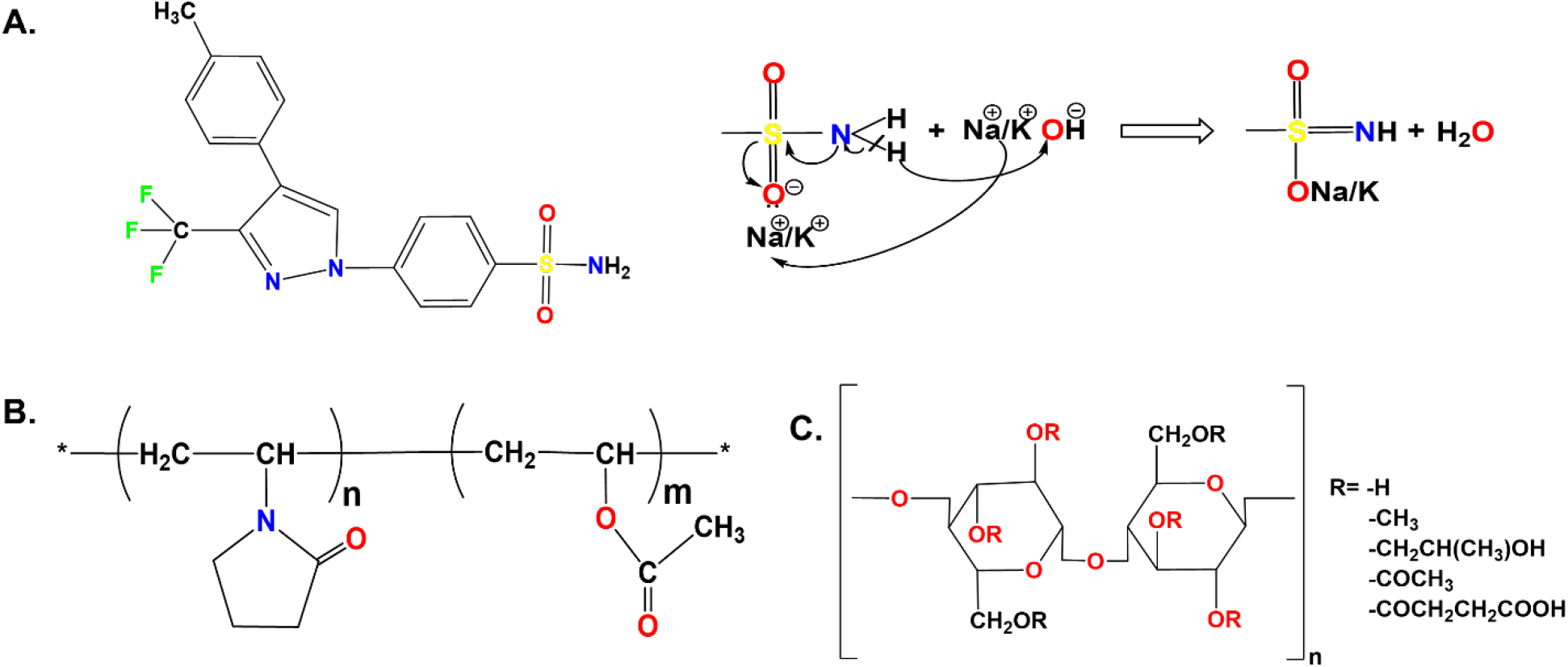
Chemical structures of CEL and the polymer excipients studied. **(A)** CEL in neutral form with scheme showing the formation of the CEL anionic salt in the presence of NaOH or KOH. **(B)** Poly-(1-vinylpyrrolidone-co-vinyl-acetate) (PVP-VA). The polymer studied, PVP-VA 64, has a ratio of 6:4 by mass for the N-vinylpyrrolidone (VP) and vinyl acetate (VA) components. (C) Hypromellose acetate succinate (HPMCAS). The M-grade contains acetyl and succinoyl groups at 7-11% and 10-14% mass fractions along with methoxyl and hydroxypropoxyl groups at 21-25% and 5-9%^[66]^.

The global CEL market was valued at 1149.8 million USD in 2020 and is expected to reach 1606.1 million USD by the end of 2026, growing at a CAGR (compound annual growth rate) of 8.7% from 2021 to 2026^[19]^. The global CEL market is mostly dominated by Pfizer, which markets CELEBREX^®^. CELEBREX^®^ contains micronized CEL particles, sodium croscarmellose as a disintegrant, and povidone K30 and sodium lauryl sulphate (SLS) as solubilizers, for enhancing the solubility and oral bioavailability of CEL^[20]^. However, SLS is an anionic surfactant that causes strong mucosal irritation and structural modification to proteins and phospholipids, leading to toxicity in humans^[21–23]^. Moreover, the positive food effect observed for CELEBREX^®^ results in varying levels of absorption of CEL. Intake of CELEBREX^®^ is recommended with food to improve absorption, but due to differences in food intake and diet, interpatient variability in the pharmacokinetic parameters can impede correct dosing^[24]^. Furthermore, treatment requires the administration of high doses of CEL, generally 200-400 mg twice daily for a prolonged period for treating serious conditions, which can lead to adverse events, such as hypersensitivity reactions (anaphylaxis), abdominal pain, nasopharyngitis and arthralgia, that hamper therapy^[25, 26]^. Thus, there is an unmet need for the development of improved dosage formulations for CEL.

We previously found that supersaturated amorphous drug-salt-polymer systems, termed amorphous salt solid dispersions (ASSD), are better than conventional binary amorphous solid dispersions (ASD) in enhancing the aqueous solubility, physical stability and bioavailability of CEL^[27]^. Specifically, ASSDs generated using *in situ* salt formation of CEL, which has a pK_a_ of 11.1 and log P of 3.5, with Na^+^ and K^+^ counterions in the matrix of a Soluplus polymer exhibited the combined effects of solubility enhancement, stabilisation through ionic interactions between drug and counterions, and intermolecular interactions between drug and polymer, that maintain the supersaturation for prolonged times in the gastrointestinal tract (GIT)^[27]^. Such prolonged effects offer the prospect of lowering the dose and for once-a-day administration that together would enhance patient compliance and reduce toxicity. These findings encouraged us to investigate in the present study whether other polymers, specifically polyvinylpyrrolidone vinyl acetate 64 (PVP-VA (6:4)) and hydroxypropyl methylcellulose acetate succinate (HPMCAS) (Fig. 1B, C), might have improved effects when used with Na^+^ and K^+^ salts of CEL in ASSDs.

It is known that intermolecular drug-polymer interactions can significantly impact the biopharmaceutical performance^[8, 9, 28]^ and physical stability^[29, 30]^ of a drug. Indeed, the general rationale for experimental screening of polymers to identify excipients for a particular drug is to identify thermodynamically miscible systems that possess adhesive interactions (specific hydrogen bonding, hydrophobic or van der Waals interactions), between the drug and the polymer in order to hinder drug-drug aggregation^[31, 32]^, and thereby prevent nucleation and crystal growth in the supersaturated solution during dissolution. The traditional methods to predict drug-polymer miscibility involve the calculation of solubility parameters and Flory-Huggins (F-H) interaction parameters^[33–35]^. We first applied these methods to compute solubility parameters to assess CEL-polymer miscibility for HPMCAS and PVP-VA. However, the results obtained did not correlate well with the measured physicochemical and biopharmaceutical attributes of the amorphous formulations. Therefore, we employed molecular dynamics (MD) simulations to evaluate drug-polymer interactions in the presence of aqueous solution.

Recent studies have employed a combination of molecular docking and MD simulations^[36]^ or MD simulations alone^[37–39]^ to investigate interactions between drugs and excipients. Moreover, in addition to atomistic MD simulations, coarse-grained MD simulations^[40, 41]^ have been carried out to understand the mechanisms of inhibition of drug-drug aggregation in the presence of an excipient. Recently, Ouyang and colleagues developed machine learning methods to aid the selection of stable drug-polymer complexes for solid dispersion formulations and their evaluation by enthalpy calculations^[42, 43]^. However, none of these studies estimated the strength of drug-drug interactions in the presence and absence of polymer, or in the presence of ions. Although molecular level mechanistic studies to elucidate the drug-polymer interplay in both solution and solid state have been performed using high-end analytical techniques^[6, 28, 32]^, the intermolecular interactions in an aqueous environment that correlate with the *in vivo* fate of a drug remain largely unexplored, but are particularly amenable to investigation by MD simulation.

Here, MD simulations of neutral and anionic forms of CEL with different polymers in aqueous solution were performed and compared with *in vitro* and *in vivo* experimental measurements for novel supersaturated ASSDs that have the potential to address the aforementioned problems of drug-drug aggregation and crystallization. We find that hindrance of drug aggregation is attributable to stronger drug-polymer interactions in the ionized drug state in ASSDs compared to the non-ionized drug in conventional ASDs. Of the systems studied, the simulations reveal the most stable intermolecular interactions between anionic CEL and PVP-VA, correlating with experimental observations which show prolonged stability along with ameliorated dissolution and pharmacokinetic profiles, offering the prospect of less frequent administration and lower doses with cost-effectiveness and fewer side effects.

## RESULTS

### Determination of drug-polymer miscibility

The drug-polymer miscibility was first calculated and then determined experimentally. The Hildebrand solubility parameter (δ) of CEL and the two polymers, PVP-VA and HPMCAS, was calculated using the Fedors and Hoftyzer-Van Krevelen (HVK) group contribution methods. The difference in the δ values of CEL and each polymer was small (<7 MPa^1/2^), indicating the possibility of miscibility of CEL in these polymers (Table 1). The difference was smaller for the CEL-HPMCAS system than for the CEL-PVP-VA system (0.65 vs 1.01 MPa^1/2^), indicating that the CEL-HPMCAS system should have a higher miscibility than the CEL-PVP-VA system. Estimation of the Flory–Huggins (F-H) interaction parameter, χ, for the two CEL-polymer systems at 25°C using the solubility parameter approach gave low positive values (Table 1). The closeness of the χ value for the CEL-HPMCAS system to zero suggests higher miscibility compared to the CEL-PVP-VA system at room temperature. However, Differential Scanning Calorimetry (DSC) thermograms for the physical mixtures (PM) of CEL with each of the two polymers revealed greater melting point depression for mixtures with PVP-VA, indicating higher miscibility with CEL of PVP-VA than HPMCAS (Fig. S1).

**Table 1.**
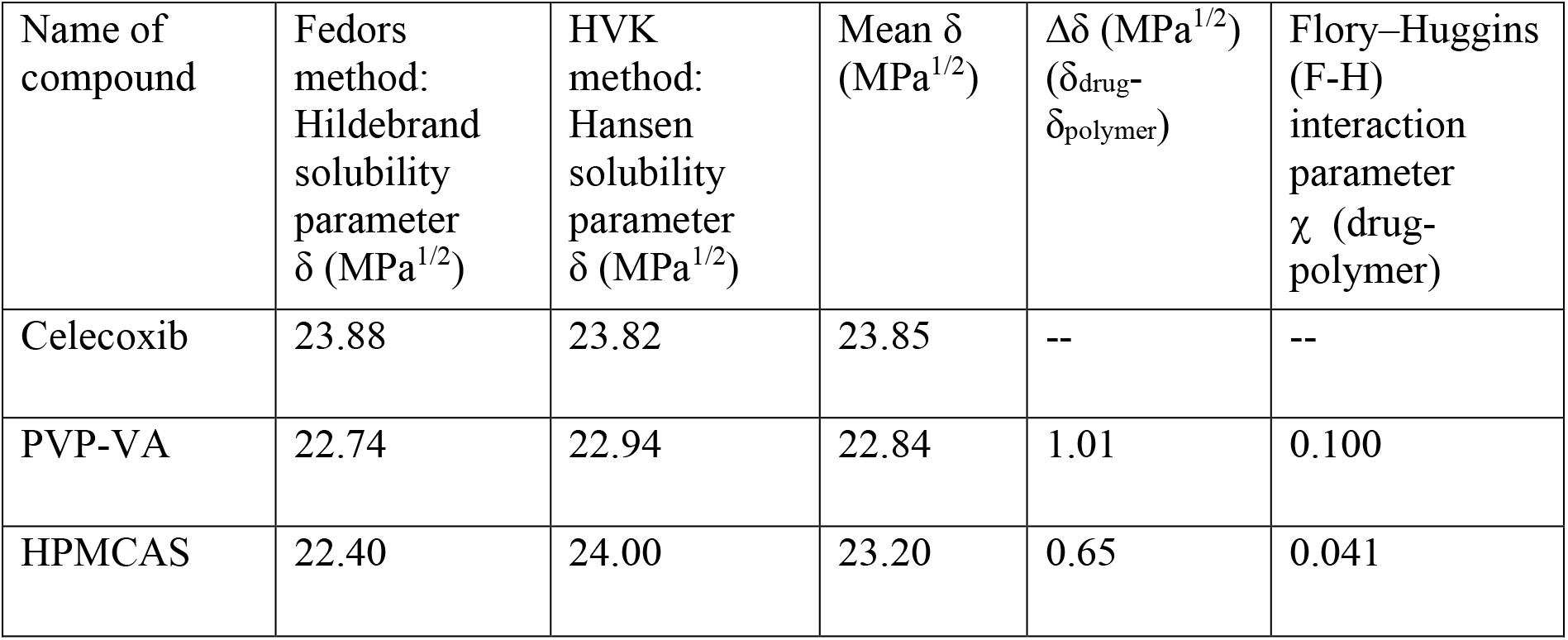
Solubility parameter values calculated by the Fedors and HVK methods for CEL and the two polymers studied.

### Preparation and characterization of formulations

Binary ASD formulations of CEL with PVP-VA or HPMCAS and ASSD formulations of CEL with Na^+^ or K^+^ and one of these two polymers were prepared using the components and procedure given in the Methods section and illustrated in Fig. S2. All the prepared formulations were white, free-flowing powdered solid dispersions. Optical light microscopy revealed long needle and plate-shaped crystals of CEL in non-polarized mode, and showed double refraction or birefringence in polarized mode (Fig. S3a), consistent with the anisotropic crystalline nature of many drugs^[44]^. All the ASD and ASSD formulations presented small spherical particles in non-polarized mode and, like amorphous CEL when analysed within few hours of preparation (Fig. S3b), no birefringence in polarized mode (Fig. S3(c-h)) due to isotropy and the absence of a periodic crystalline lattice, revealing their amorphous nature and indicating the role of the polymers in stabilizing the high energy amorphous form of CEL by slowing or preventing its recrystallization during storage, as evident from birefringence in microscopy (Fig S12b), and at higher temperature as observed in the DSC thermogram (Fig. S4B).

These results were further supported by DSC and Powder X-ray Diffraction (PXRD). In DSC, a sharp melting endotherm was observed at 163.6°C for crystalline CEL (Fig. S4A), whereas the polymers lacked a sharp melting endotherm and instead exhibited a glass transition temperature (T_g_) (Fig. S4B), indicating their amorphous nature. Moreover, the observation of T_g_ values in the thermograms of all of the drug-polymer formulations confirmed their amorphous nature (Fig. S4A). These T_g_ values range from 71.5 to 108.8 °C and are higher than the T_g_ of pure amorphous CEL (57.9 °C), indicating better stability of the formulations^]^. Notably, amorphous CEL showed recrystallisation at high temperature followed by a melting peak at 164.9°C in the DSC thermogram (Fig S4B) whereas no recrystallisation was observed for the drug-polymer formulations, thus confirming their stability in amorphous form. The T_g_ values of the ASSDs are higher than those for the ASDs and are higher for the K^+^-containing ASSDs than the Na^+^-containing ASSDs, with the highest value being observed for the CEL-K-PVP-VA ASSD. The higher T_g_ values for the ASSD formulations indicate strong ionic interactions between CEL and the counterions and they form more stable amorphous systems. While the smaller Na^+^ ion might be expected to interact more strongly with CEL than the larger K^+^ ion, the K^+^ ion has a lower ionization enthalpy and may therefore have a greater tendency to form ionic interactions with CEL in the amorphous form.

In PXRD, CEL showed the characteristic peak pattern of crystalline polymorphic form III whereas a halo pattern was observed for amorphous CEL, both polymers, and the ASDs and ASSDs, confirming their amorphous nature (Fig. S5).

### Determination of intermolecular interactions

Fourier transform infrared spectroscopy (FTIR), micro-Raman spectroscopy and nuclear magnetic resonance (NMR) spectroscopy were performed to examine the molecular interactions between CEL, in both neutral and ionic states, and the polymers in supersaturated formulations.

#### FTIR spectroscopy

The overlay of FTIR spectra revealed that in the amorphous formulations, the medium intensity doublet bands, corresponding to the N–H stretching of the –NH_2_ group, were diffused and broadened, and that the asymmetric stretching vibration of the –SO_2_ group of CEL shifted to a lower frequency in the CEL-PVP-VA ASD and disappeared in both the CEL-PVP-VA ASSDs whereas the peaks persisted in amorphous CEL (Fig. 2A). This indicates the participation of the –NH_2_ group of CEL in intermolecular hydrogen bonding with the polymer^[45]^ and the –SO_2_ group of CEL in salt formation with the counterion due to the greater electronegativity of the oxygen atom than the nitrogen atom, which is attributed to the -R effect (negative resonance) shown by the sulphonamide group (Fig. 1A). The interaction of CEL with the counterion can also be observed in the crystal structure of the CEL-Na^+^ salt^[46]^. The weaker electrostatic interaction between CEL and the K^+^ counterion is due to the larger ionic radius and lower charge density of K^+^ as compared to the Na^+^ counterion^[27, 47]^.

**Fig. 2.**
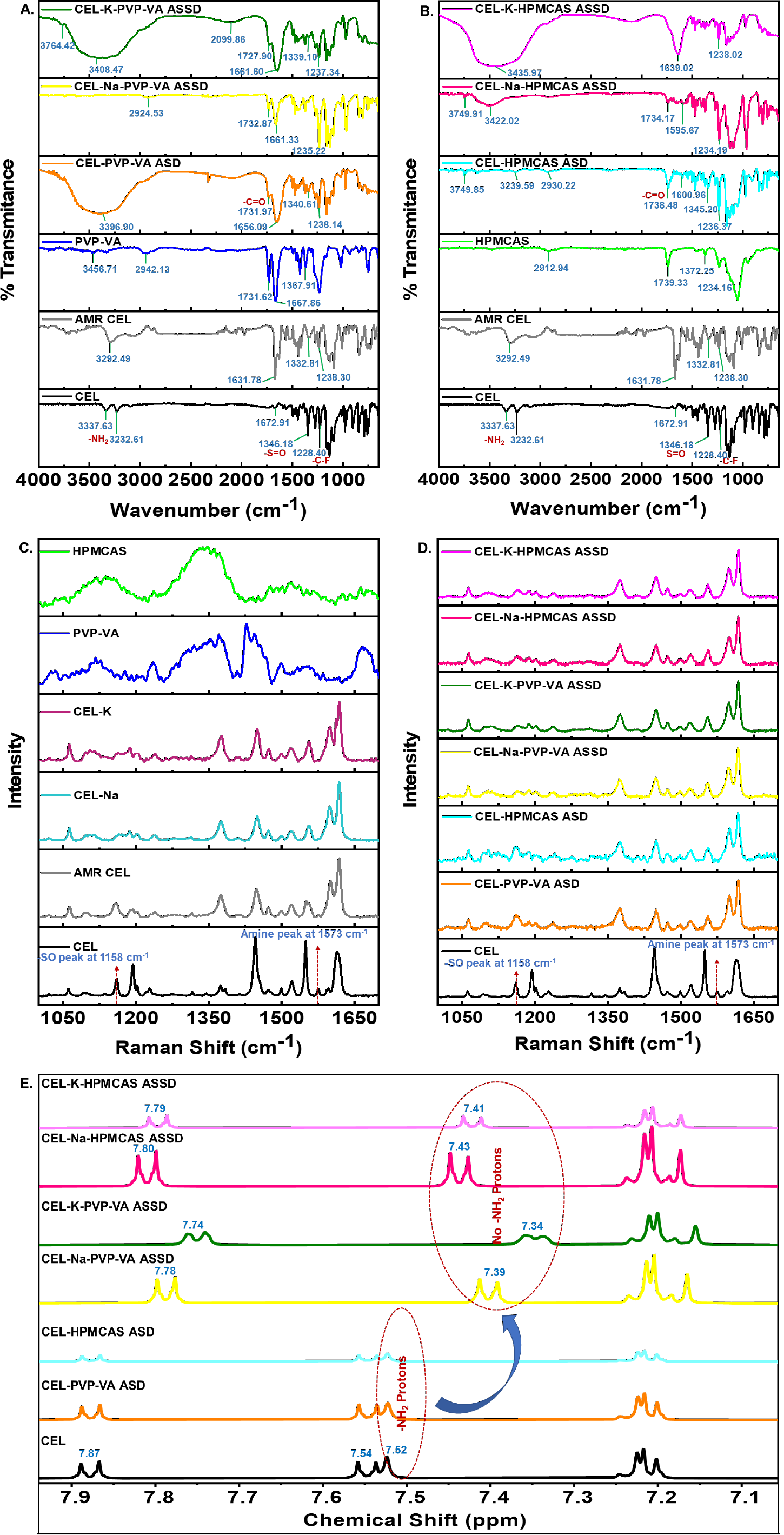
Spectroscopic techniques reveal intermolecular interactions between CEL and the polymer in amorphous formulations. **(A,B)** FTIR-ATR spectra of crystalline and amorphous (AMR) CEL, and the PVP-VA (A) and HPMCAS (B) polymers and CEL-polymer formulations. The spectra show the disappearance of the -S=O peak in ASSDs, indicating the involvement of the -S=O group in salt formation (Figure 1), and the broadening of the -NH stretching peak of the -NH_2_ group in the ASDs and ASSDs. **(C,D)** Micro-Raman spectra of crystalline (C) CEL and its ASD and ASSD formulations, and (D) CEL, AMR CEL, CEL Na^+^/K^+^ salts, and the PVP-VA and HPMCAS polymers. The spectra for the ASSD formulations and the CEL Na^+^/K^+^ salts show the disappearance of the -S=O peak, indicating salt formation, and the disappearance of the -NH stretching peak in both ASD and ASSD formulations, revealing CEL-polymer hydrogen-bonding. **(E)** NMR spectra between 7.1 and 7.9 ppm of CEL and its ASD and ASSD formulations. The complete spectral region is shown in Fig. S6. The spectra show the upfield shift in the aromatic protons of CEL at 7.87 and 7.54 ppm and the disappearance of the - NH_2_ proton peak at 7.52 ppm in the ASSD formulations, indicating salt formation.

The –C=O stretching frequency of the VA moiety of PVP-VA at 1667 cm^-1^ shifted downfield in the ASD and ASSDs, suggesting the formation of a hydrogen bond between the –C=O group of the polymer and the –NH group of CEL. The –C=O stretching frequency of the VP moiety of PVP-VA at 1731 cm^-1^ shifted in the ASSDs but there was no change in frequency for the ASD. Moreover, the –C–F symmetric stretching vibrations peak at 1228 cm^-1^ in CEL shifted to higher frequencies in all the PVP-VA and HPMCAS formulations and its intensity reduced, confirming the formation of disordered amorphous structures^[48]^ (Fig. 2A,B).

For the CEL-HPMCAS systems, the doublet of –NH_2_ protons of CEL at 3337 cm^-1^ and 3232 cm^-1^ shifted and broadened in all the formulations, indicating the formation of a hydrogen bond whereas the peaks still persisted in amorphous CEL and shifted slightly to 3292.49 cm^-1^. The –S=O peak at 1346 cm^-1^ in CEL shifted to 1332.81 cm^-1^ in amorphous CEL and 1345 cm^-1^ in the ASD, but dissipated in both CEL-HPMCAS ASSDs, indicating salt formation. The –C=O stretching frequency of HPMCAS at 1739 cm^-1^ shifted to 1738 cm^-1^ in the ASD and 1734 cm^-1^ in the CEL-Na-HPMCAS ASSD, but disappeared in the CEL-K-HPMCAS ASSD. Also, the broad –OH stretching peak at 2912 in HPMCAS get shifted to 2930 in CEL-HPMCAS ASD, but vanished in both CEL-HPMCAS ASSDs, suggesting hydrogen bond formation between the –NH group of CEL and the –C=O or – OH groups of the polymer (Fig. 2B).

These results indicate that the counterion (Na^+/^K^+^) interacts with CEL via strong ionic interactions resulting in salt formation. Also, the *in situ* generated salt and variations in the peak shifts suggest that CEL is hydrogen-bonded to both the –C=O groups of the PVP-VA polymer. These hydrogen bonds may help to maintain the supersaturation of CEL in solution for long times by inhibition of nucleation and crystal growth. However, the interactions are more evident from Raman spectroscopy in Fig 2C,D.

#### Micro-Raman spectroscopy

We previously described the characteristic peaks of CEL present in the micro-Raman spectrum range of 1000-1700 cm^-1^ ^[27]^ and here we focused on the same range (Fig. 2C,D). Crystalline and amorphous CEL exhibited a characteristic peak of –S=O asymmetric stretching at 1158 cm^-1^ which was also observed in both CEL PVP-VA and HPMCAS ASDs. However, the disappearance of this peak in the CEL (Na^+^/K^+^) salts and all the CEL ASSDs confirms electrostatic interactions between the CEL –S=O group and the counterions (Na^+^/K^+^), resulting in salt formation. Furthermore, the – NH peak present for crystalline CEL at 1573 cm^-1^ was of low intensity in amorphous CEL and disappeared in all the formulations, confirming the hydrogen bonding interactions between the –NH group of CEL and the –C=O group of PVP-VA polymer and, –C=O or – OH groups of HPMCAS polymer as observed in FTIR spectra. Thus, the FTIR and micro-Raman spectroscopy both reveal *in situ* salt formation of CEL with the counterion and CEL hydrogen bonding with the polymer.

#### NMR spectroscopy

Chemical shifts in the aliphatic and aromatic regions of CEL as a result of alterations in the proton microenvironments were measured by NMR^[32]^ for CEL and its amorphous formulations (Fig. S6, Fig 2E). The upfield shift in the two doublet peaks of aromatic protons attached to the sulphonamide group of CEL and the disappearance of the –NH_2_ peak at a chemical shift of 7.52 ppm in ASSDs but not ASDs (Fig. 2E) confirmed salt formation between CEL and the counterion (Na^+^/K^+^) due to loss of a proton from the –SO_2_NH_2_ group, leading to shielding of the neighbouring aromatic protons in CEL. In the –SO_2_NH_2_ group, the −NH_2_ and −SO_2_ act as proton donor and acceptor, respectively. The −SO_2_, being a strong electron-withdrawing group, will undergo a -R effect due to conjugation between the lone pairs of electrons of the oxygen atoms and the pi electron of the resonating system. There is a delocalisation of pi electrons towards the oxygen atoms, which further extracts electrons from the N atom of the –NH_2_ group and leads to formation of the -S=NH bond. This effect will lead to more electron-rich oxygen atoms that can make an ionic interaction with the positively charged counterion (Fig. 1A). Furthermore, the upfield shifts of the aromatic protons in the ASSDs indicate a change in their electron density because of drug-polymer interactions. The doublet peaks of the aromatic protons in CEL at chemical shift values of 7.87 and 7.54 ppm were shifted upfield in the ASSDs, whereas no such alteration in the peaks was observed for the ASD formulations (Fig. 2E). Hydrophilic polymers can interact with drug molecules by forming electrostatic interactions or by hydrogen bonding, thereby inhibiting drug-drug aggregation and maintaining supersaturation for prolonged times^[27]^. The CEL chemical shifts, along with the enhanced T_g_ obtained in the DSC studies, further confirm the *in situ* salt formation between CEL and the counterions (Na^+^/K^+^) and the drug-polymer interactions in the ASSD formulations.

### Supersaturation and precipitation studies

The extent of CEL supersaturation in the ASSDs and ASDs was studied at 37 °C using an ultraviolet (UV) spectrophotometer. CEL and all CEL-polymer formulations were dissolved separately in methanol at 5 mg/mL CEL concentration and poured into 25 mL water. Supersaturation experiments were conducted in water because water can act as a strong plasticizer for amorphous systems and cause them to recrystallize^[49]^. Initially, a high absorbance was observed for CEL followed by a rapid decline due to precipitation of CEL in water. The CEL-K-PVP-VA ASSD generated maximum supersaturation of CEL for 4 h due to the hydrophilic nature of the PVP-VA polymer, however the CEL-Na-PVP-VA ASSD showed a slower decline in CEL concentration for 40 min and maintained it for 4 h (Fig. S7b). This confirmed the greater hindrance of water-induced crystallization of CEL in the CEL-K-PVP-VA ASSD than in the CEL-Na-PVP-VA ASSD. CEL in the binary ASDs with both the polymers showed reduced absorbance compared to the ASSDs due to crystallization from its supersaturated state, which can be attributed to weaker intermolecular interactions between CEL and the polymer in the neutral state than in the ionic state. The ASSDs with HPMCAS were less performant in the maintenance of CEL supersaturation (Fig. S7a) due to the lower aqueous solubility of HPMCAS. Due to the presence of the comparatively hydrophobic methoxy and acetate substituents, HPMCAS is water-insoluble when unionized at pH values below 5 and forms a colloidal solution at the intestinal pH of 6.0–7.5. This ultimately results in poor aqueous solubility of the polymer even in its ionized state at the small intestinal pH and leads to the formation of colloidal polymer aggregates in aqueous solutions that promote interactions within the polymer^[50]^. The better behaviour of the systems containing PVP-VA than those containing HPMCAS is contrary to the predictions from the calculation of drug-polymer miscibility parameters (Table 1) but correlates well with the observed melting point depression in DSC thermograms which indicated high CEL-polymer miscibility in the case of PVP-VA (Fig. S1A). Therefore, to investigate the physical mechanisms underlying the supersaturation of CEL, we next performed molecular modelling and simulation.

### *In silico* molecular modelling

The systems simulated, labelled a-j, are listed in Table 2.

**Table 2.** Specifications of the simulated systems to study the aggregation of CEL. MD simulations were carried out for polymeric excipients without CEL (a,b), and for the neutral (c,d,g,i) and anionic (e,f,h,j) forms of CEL in the absence (c-f) and presence (g-j) of a polymeric excipient. Each drug-polymer system was simulated in duplicate, generating trajectories referred to as replica-1 and replica-2. For the systems without polymer, two systems with different initial box sizes were each simulated once.

| System | # of CEL molecules and ionization state | Polymer and its length (Å)<sup>#</sup> | Periodic box size (Å)<sup>*</sup> | # of water molecules | # of Na<sup>+</sup> ions | # of Cl<sup>-</sup> ions | # of replica simulations | Simulation length (ns) |
| --- | --- | --- | --- | --- | --- | --- | --- | --- |
| a | 0 | PVP-VA (75.1) | X=154.2;<br>Y= 94.3;<br>Z= 99.9 | 46569 | 0 | 0 | 1 | 66 |
| b | 0 | HPMCAS (79.2) | X =158.2;<br>Y = 127.3;<br>Z = 99.2 | 58720 | 4 | 0 | 1 | 66 |
| c | 13 neutral | --- | 112 | 138672 | 9 | 9 | 1 | 516 |
| d | 11 neutral | --- | 106 | 117072 | 7 | 7 | 1 | 456 |
| e | 13 anionic | --- | 112 | 138672 | 22 | 9 | 1 | 916 |
| f | 11 anionic | --- | 106 | 117066 | 18 | 7 | 1 | 1042 |
| g | 13 neutral | PVP-VA (43.9) | 112 | 46031 | 9 | 9 | 3 | 356/356/556 |
| h | 13 anionic | PVP-VA (43.9) | 112 | 46038 | 22 | 9 | 2 | 356 |
| i | 11 neutral | HPMCAS (72.2) | 106 | 38819 | 11 | 7 | 2 | 376 |
| j | 11 anionic | HPMCAS (72.2) | 106 | 38831 | 22 | 7 | 2 | 376 |

<sup>#</sup>The polymer length is given at the beginning of the respective simulation(s).
<sup>\*</sup>The initial box was cubic unless specified by giving the X, Y and Z dimensions.

#### Simulation of polymers

Explicit solvent MD simulations of the individual polymers were initiated with either a PVP-VA or a HPMCAS oligomer in a rather extended modelled conformation in a periodic box of aqueous solution (Fig. 3). The oligomers of the two polymers deviated from linearity and adopted bent structures during the simulations. The all-atom root-mean-square deviation (RMSD) values showed convergence during the simulations (Fig. S8 a and b).

**Fig. 3.**
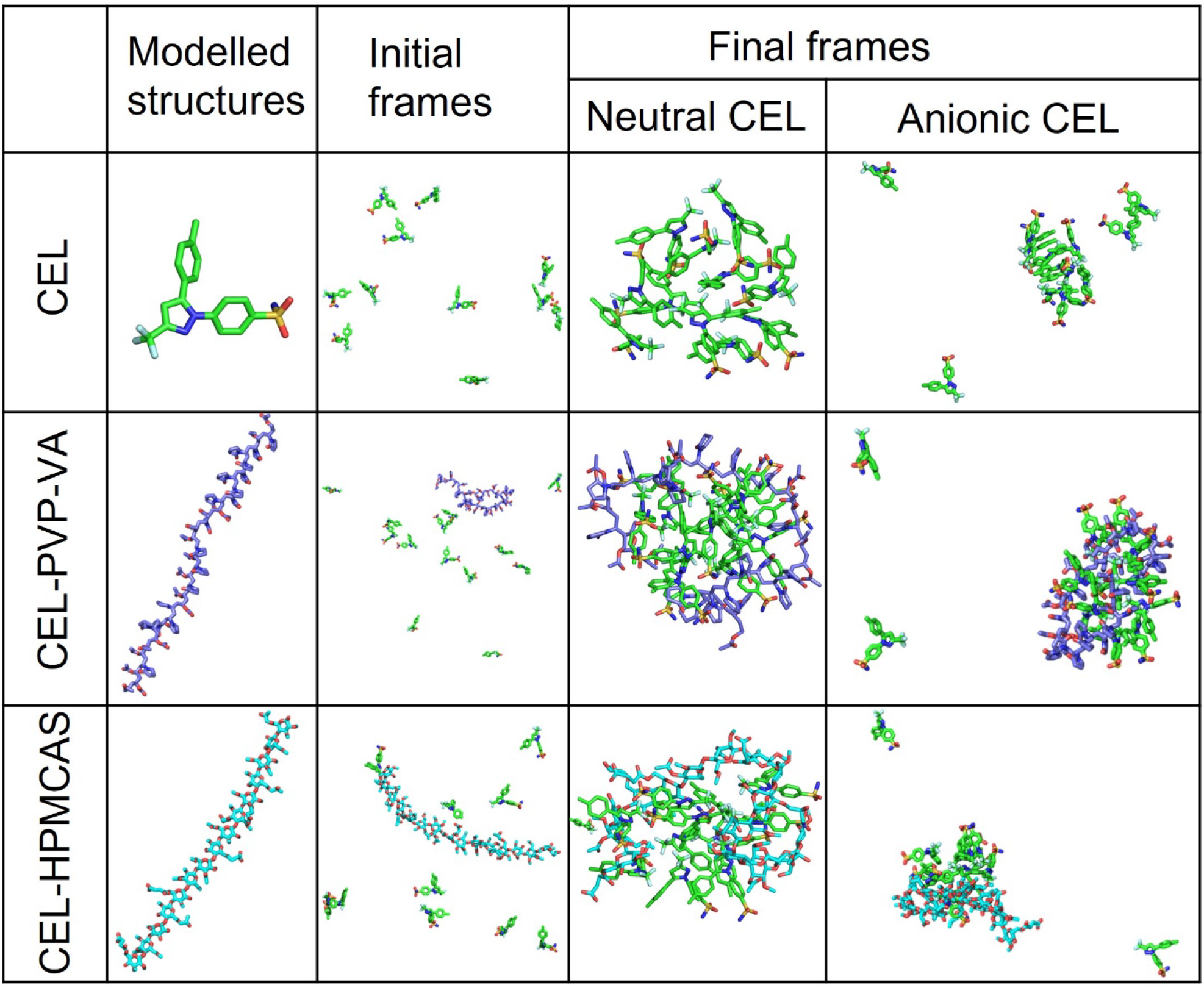
Modelled structures of the molecules simulated and snapshots of initial and final frames of the simulated systems of CEL alone and CEL with either PVP-VA or HPMCAS. Initially, the CEL molecules were placed in random positions and orientations in a periodic box of water molecules and ions without or with an oligomer. The final frames for the simulation times given in Table 2 are shown for simulation replica-1 for systems with neutral or anionic forms of CEL. For the neutral form of CEL, aggregation of the CEL molecules alone or with the oligomer is observed during all the simulations. For the anionic form of CEL, a lower aggregation propensity was observed with some CEL molecules remaining free during the simulations (the final snapshot after simulation for 916 ns for anionic CEL without oligomer is shown for system e but in system f, which also lacked oligomer (Table 2), no aggregation of anionic CEL molecules was observed in a simulation of 1042 ns duration).

#### Simulation of CEL aggregation

Simulations were conducted to model the amorphous forms of CEL (both neutral and anionic forms) in the absence of polymer and the presence of ∼0.01M NaCl (see Table 2 and Methods). CEL aggregates during simulations in aqueous solution at supersaturated concentration. The neutral form of CEL aggregated within a few hundred ns (Fig. 3). On the other hand, the anionic form of CEL only partially aggregated during one simulation (system e, final snapshot at 916 ns shown in Fig. 3) and remained dissociated during the other simulation (system f simulated for 1042 ns). In the aggregates of CEL, no specific non-covalent interactions were found, except that for neutral celecoxib, an intermolecular hydrogen bonding interaction was observed between the sulfonamide -NH_2_ group of one CEL molecule and the -SO_2_ group of another molecule (see radial distribution function (RDF) in Fig. 4A (a)). The same interaction was much reduced for the anionic form of CEL (Fig. 4A (b)). These results support the faster aggregation of CEL in the neutral state than in the ionic state, in the absence of polymer, due to stronger interactions between the CEL molecules.

**Fig. 4.**
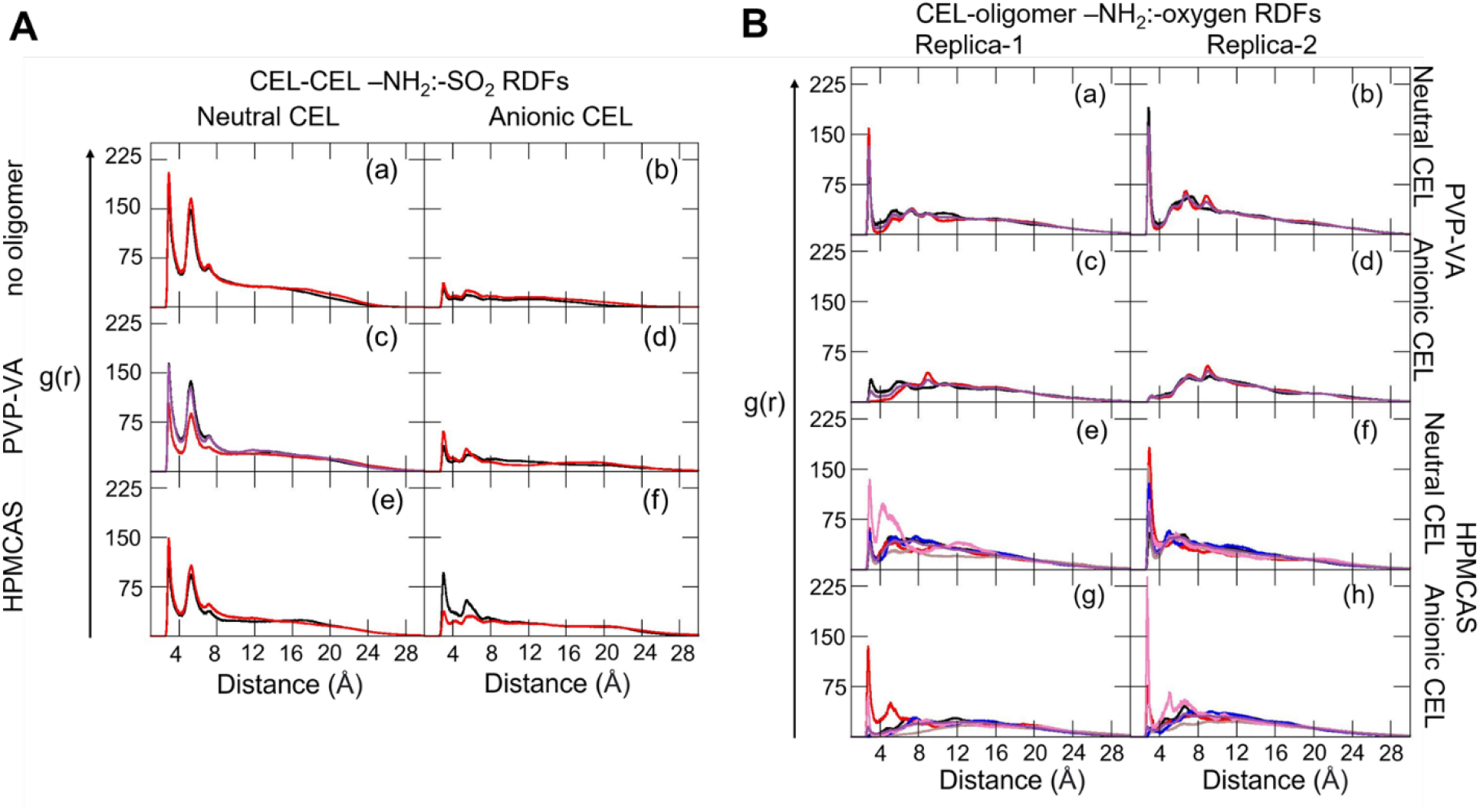
Radial distribution functions (RDFs) computed from all-atom explicit solvent MD simulations showing CEL-CEL and CEL-oligomer intermolecular interactions. **(A).** Intermolecular CEL:CEL RDFs between the sulfonamide -NH_2_ group of one CEL molecule and the SO_2_ group of another (replica-1: black, replica-2: red, replica-3: purple) for the neutral (left) and anionic (right) forms of CEL without polymer (a,b), with PVP-VA (c,d), and with HPMCAS (e,f). The RDFs show reduced neutral CEL-CEL interactions in the presence of polymer and tend to zero with increasing distance because the molecules aggregate. **(B).** Intermolecular CEL:oligomer RDFs computed from the replica-1 (left) and replica-2 (right) MD simulations (see Fig. S9B for replica-3). RDFs are shown for: the sulfonamide -NH_2_ group of the neutral (a,b) and anionic (c,d) forms of 13 CEL molecules and the carbonyl oxygen atoms of vinyl acetate (black) and vinylpyrrolidone (red) moieties and all oxygen atoms (purple) of the PVP-VA oligomer; sulfonamide -NH_2_ group of neutral (e,f) and anionic (g,h) forms of 11 CEL molecules and oxygen atoms of methoxy (black), succinoyl (brown), acetyl (blue), hydroxypropoxy (red), hydroxy (pink) and all oxygen atoms (purple) of the HPMCAS oligomer.

#### Simulation of CEL-polymer systems

Simulations were next performed for CEL in the presence of polymer and ∼0.01M NaCl and the aggregation propensity of the CEL:polymer systems was compared with that of the CEL-CEL systems (see Table 2 and Methods). The CEL-polymer interactions modulate the CEL:CEL hydrogen-bonding interactions. We performed two sets of MD simulations per CEL-polymer combination and observed formation of drug-polymer aggregates showing structural convergence after about half the simulation length (see Fig. S8). We therefore and analysed the converged parts of the trajectories in terms of hydrogen bond contact occupancy and RDFs, and computed interaction free energies with the MM/GBSA method. To reduce the higher statistical uncertainty in the computed free interaction energies for the neutral CEL:PVP-VA system compared to the other simulated systems, we carried out a third replica simulation for a longer time for this system. The third replica showed similar convergence as regards RMSD within the first 150ns and this was maintained up to the end of this simulation at 556 ns (Fig S8e). The radius of gyration of the oligomer chain also converged to a similar value to that observed for the first two replicas (see Fig. S8i). The MM/GBSA energies computed for time intervals to 356 ns or to 556 ns were very similar (see Table S1 and S2), indicating that the replica simulations of this system up to 356 ns were of sufficient duration to investigate the properties of the neutral CEL-PVP-VA aggregates.

The computed RDFs are shown in Figure 4. Differences in g(r) peak heights between replicas reflect the transient and rather non-specific nature of hydrogen-bonding interactions between the sulfonamide nitrogen and oxygen atoms of two CEL molecules, or between CEL and oligomer atoms, as apparent from visual inspection of the trajectories. For example, for the negatively charged celecoxib and HPMCAS system (**Fig. 4B** g-h), we observed that a hydrogen-bonding interaction between the nitrogen atom of the sulfonamide group in CEL and a hydroxyl group in HPMCAS was present throughout the trajectories in both replica simulations. However, this hydrogen-bonding interaction did not occur between a specific pair of CEL and HPMCAS hydroxyl atoms. Rather, it occurred between any of the eleven CEL molecules and the HPMCAS hydroxyl atoms. Moreover, the RDFs were computed by generating histograms of the number of particles found as a function of distance and normalizing by the expected number of particles at the corresponding distances. Since the modelled HPMCAS chain has eight hydroxyl groups in hydroxypropyl moieties but only three hydroxyl groups attached to the tetrahydropyran rings, the discrepancy in the RDF peak heights in the two replicas shown in Fig. 4B g and h can be attributed, in part, to different normalization factors.

##### Interactions between

*PVP-VA and CEL.* For PVP-VA, a hydrogen bond (average occupancy among two sets ∼43%) was observed between the sulfonamide −NH_2_ group of CEL and the carbonyl (−C=O) group on the VP and VA moieties of PVP-VA. The RDF between the −NH_2_ group of the neutral form of CEL and all oxygen atoms of the PVP-VA polymer shows a peak (Fig. 4B (a-b), S9B) at 2.8±0.01 Å, whereas this peak is absent for the anionic form of CEL. In the presence of PVP-VA, the relative g(r) value at the peaks in the RDFs of the neutral CEL sulfonamide −NH_2_ and −SO_2_ groups is lower (Fig. 4A (c)) than in the absence of polymer (Fig. 4A (a)) for all replicas. Thus, the CEL:CEL hydrogen-bonding interactions are weakened by the presence of the PVP-VA oligomer.

##### Interactions between HPMCAS and CEL

The RDF was computed for each of the five individual functional groups of HPMCAS and for all the five groups together. The g(r) values between the neutral form of CEL and the hydroxypropoxy and hydroxyl groups of HPMCAS are the highest in both sets of simulations (Fig. 4B (e-f)). The g(r) at a distance ≤3 Å between the hydroxypropoxy group of HPMCAS and the anionic form CEL (Fig. 4B (g-h)) is less than for the neutral form of CEL, but much higher than the carbonyl oxygen of the VA group of PVP-VA (Fig. 4B (c-d)). As observed for PVP-VA, the relative g(r) value of the RDF peaks for the sulfonamide −NH2 and −SO2 groups of neutral CEL in the presence of HPMCAS, is decreased (Fig. 4A (e)) compared to in the absence of polymer (Fig. 4A (a)) in both replicas. Thus the CEL:CEL hydrogen-bonding interactions are also weakened by the presence of the HPMCAS oligomer.

##### Effect of polymers on CEL aggregation

Upon analysing the MD trajectories, we found stable CEL:oligomer complexes and a small reduction in the magnitude of the CEL:CEL interaction energy in the presence of an oligomer for the neutral form of CEL (Fig. 5 and Table S1). For the anionic form of CEL, partial or no aggregation was observed in the absence of oligomer, whereas relatively stable CEL aggregates were observed in the presence of oligomers. Of the two polymers considered here, PVP-VA formed more energetically stable complexes with CEL. Interestingly, the PVP-VA oligomer tended to form a relatively more stable complex with the anionic than the neutral form of CEL. This can be attributed to the higher degree of intermolecular contact between the fluorine atoms of anionic CEL and the carbonyl carbon atoms of the PVP-VA oligomer as observed in the RDFs between these atoms (Fig. S9). This interaction strengthened the electrostatic and van der Waals forces between the PVP-VA oligomer and the anionic form of CEL (Table S2). On the other hand, the electrostatic interaction energy between the anionic form of CEL and the HPMCAS oligomer is unfavorable due to repulsive interactions between the negatively charged sulfonamide group of CEL and the oxygen atoms of the oligomer. Nevertheless, the overall energy of the complex is negative due to the favorable van der Waals and solvation energies.

**Fig. 5.**
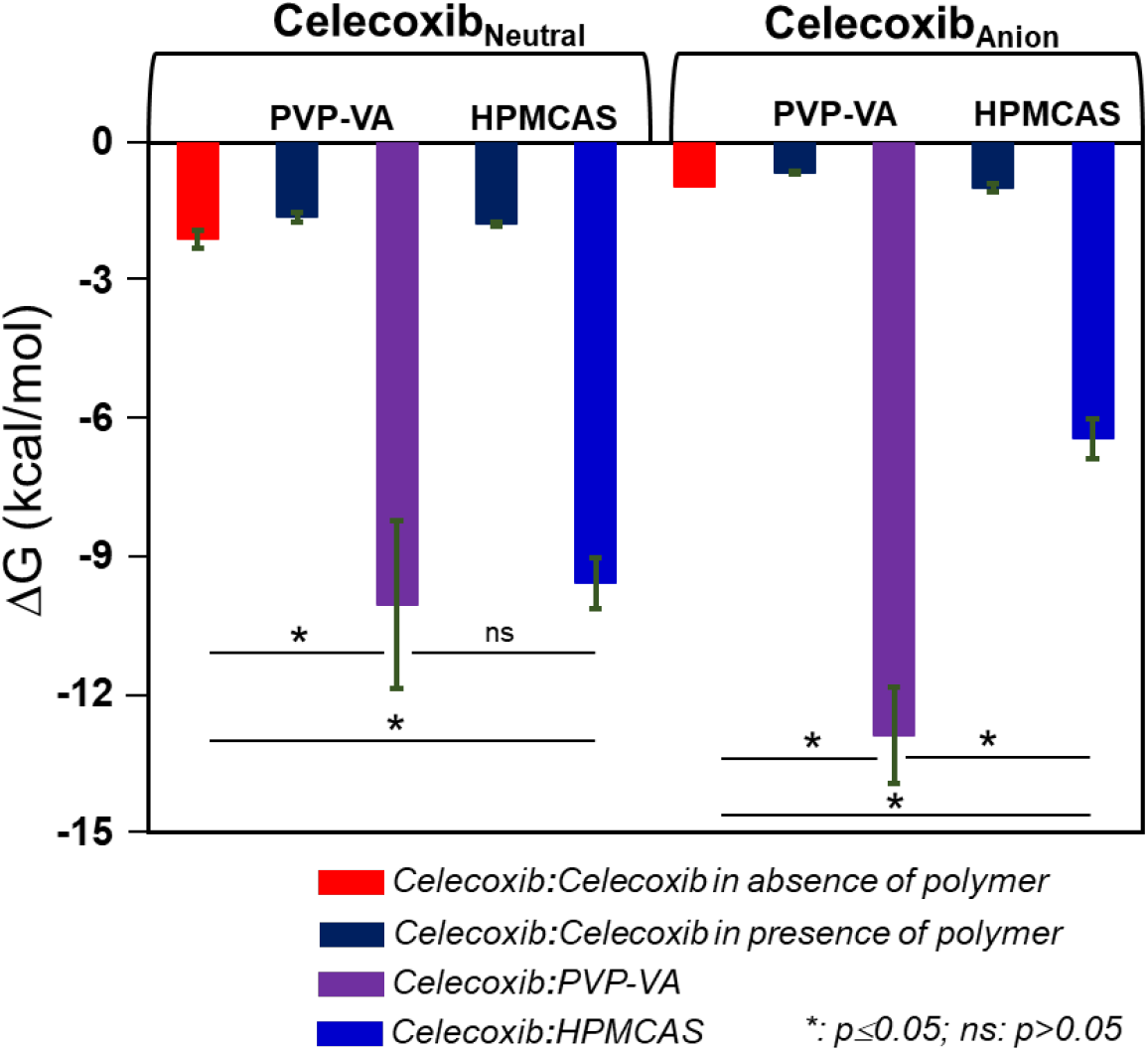
Computed interaction free energies for the drug and drug-polymer systems indicate suitable polymer excipients. Computed MM/GBSA energies (kcal/mol) for the CEL:CEL (red, dark blue), CEL:PVP-VA (purple) and CEL:HPMCAS (blue) interactions in the aggregates formed in the MD simulations show more energetically favorable interactions of PVP-VA with CEL than HPMCAS, particularly for the anionic form of CEL, with a corresponding tendency to energetically weaker CEL:CEL interactions. These trends are consistent with the better excipient properties of PVP-VA than HPMCAS for the CEL ASSD. The average interaction free energies and their standard deviations computed from snapshots at 100 ps intervals from the equilibrated parts of the replica simulations for each system are shown along with p-value ranges for relevant interaction free energy differences. Computed values are given in Table S1, their decomposition is given in Table S2, and the results of bootstrapping of the computed energies are given in Table S3.

We found that as the strength of oligomer:CEL interaction energy increases, the magnitude of the CEL:CEL interaction energy in the CEL aggregate tends to decrease. The strength of the oligomer:CEL interaction is several-fold higher per CEL molecule than that of the CEL:CEL interaction in the CEL aggregate for both neutral and anionic forms of CEL (Fig. 5). In energy component analysis, in the neutral form of the CEL aggregate, the intermolecular van der Waals component is more favourable than the electrostatic energy. For the anionic form of CEL in the presence of HPMCAS, the electrostatic energy term is highly unfavourable. However, due to the favourable solvation and van der Waals interaction energies, the overall binding free energy is favourable between CEL molecules in the anionic form and the HPMCAS oligomer. Since the CEL:PVP-VA interactions per CEL molecule are stronger than the CEL:HPMCAS interactions, for the anionic form of CEL, PVP-VA could serve as a candidate polymer for CEL for preventing its precipitation and maintenance of supersaturation for prolonged times due to more favourable intermolecular interactions between the drug and the polymer. Notably, the relative ranking of the MM/GBSA CEL:CEL and CEL:polymer binding free energies for both neutral and anionic forms of CEL for the two polymers, is opposite from that derived from the computed solubility parameters (Table 1). However, the ranking from the MD simulations, which indicates better miscibility of CEL with PVP-VA than with HPMCAS, correlates with the experimentally observed intermolecular interactions and biopharmaceutical performance as measured in the next section.

### Solubility, dissolution, and biopharmaceutical performance of the amorphous formulations

#### Apparent solubility studies

Apparent solubility studies performed in distilled water for 6 h showed different solubilities of CEL in its different formulations (Fig. 6A). Water was used as media because it can act as a strong plasticizer for amorphous systems and cause them to recrystallize^[49]^. Therefore, to check the role of amorphous formulations, both ASDs and ASSDs, in enhancing the aqueous solubility and inhibition of crystallization of CEL, experiments were performed in water. Crystalline CEL attained the maximum solubility of 4.53 μg/mL after 30 minutes, whereas for the CEL-PVPVA and CEL-HPMCAS ASD formulations, the highest solubility obtained was 34.96 and 20.26 μg/mL, respectively after 5 minutes. For the CEL-Na-PVP-VA and CEL-K-PVP-VA ASSD formulations, the maximum solubility was attained within 5 minutes and was 736.87 and 392.48 μg/mL, respectively, whereas for the CEL-Na-HPMCAS and CEL-K-HPMCAS ASSD formulations, the maximum solubility was 73.88 and 193.19 μg/mL after 15 and 30 minutes, respectively. However, the drug concentration declined afterwards for the ASSDs, whereas no such changes were observed for the ASDs. The supersaturation was maintained for the ASSDs and the concentrations achieved after 6 hours were 139.54 and 154.72 μg/mL for CEL-Na-PVP-VA and CEL-K-PVP-VA ASSDs, respectively, and 25.54 and 126.80 μg/mL for CEL-Na-HPMCAS and CEL-K-HPMCAS ASSDs, respectively. The enhancement in solubility with respect to crystalline CEL was over 40-fold in the PVP-VA ASSDs and up to 36-fold in the CEL-K-HPMCAS ASSD. The CEL-Na-PVP-VA ASSD showed higher solubility than CEL-K-PVP-VA ASSD in water at short times and then the solubility fell to about the same level as that for the CEL-K-PVP-VA ASSD for the rest of the time monitored. This might be explained by the lower T_g_ value of the former, leading to more drug release and higher solubility due to the hydrophilic nature of PVP-VA. This was observed recently for PVP-VA and a different drug compound^[51]^. In contrast, the CEL-K-HPMCAS ASSD showed higher solubility than the CEL-Na-HPMCAS ASSD throughout the measurement time despite having a slightly higher T_g_ value. This is because the comparatively hydrophobic nature of HPMCAS causes slower or low salt release in water but the release of the larger K^+^ ions, with higher aqueous solubility than Na^+^, has a greater structure-breaking effect on the water^[47]^. The overall higher solubility for K^+^-containing salt formulations is attributed to the higher polarizability of the K^+^ ion than Na^+^ ion^[52]^. The CEL embedded in the formulations is in amorphous form, as supported by DSC and XRD spectra (see Fig. S4 and S5, respectively), and dissolved immediately in the aqueous environment (spring effect) and attained supersaturation (parachute effect). The combined effect of the loss of the crystal lattice energy and the pH buffering effect of the salt in ASSD formulations accounted for the higher apparent solubility of CEL, and extended supersaturation (see Fig. S7) due to stable intermolecular interactions. This spring and parachute phenomenon is important for significant dissolution and augmented the *in vivo* bioavailability of the drug^[8]^.

**Fig. 6.**
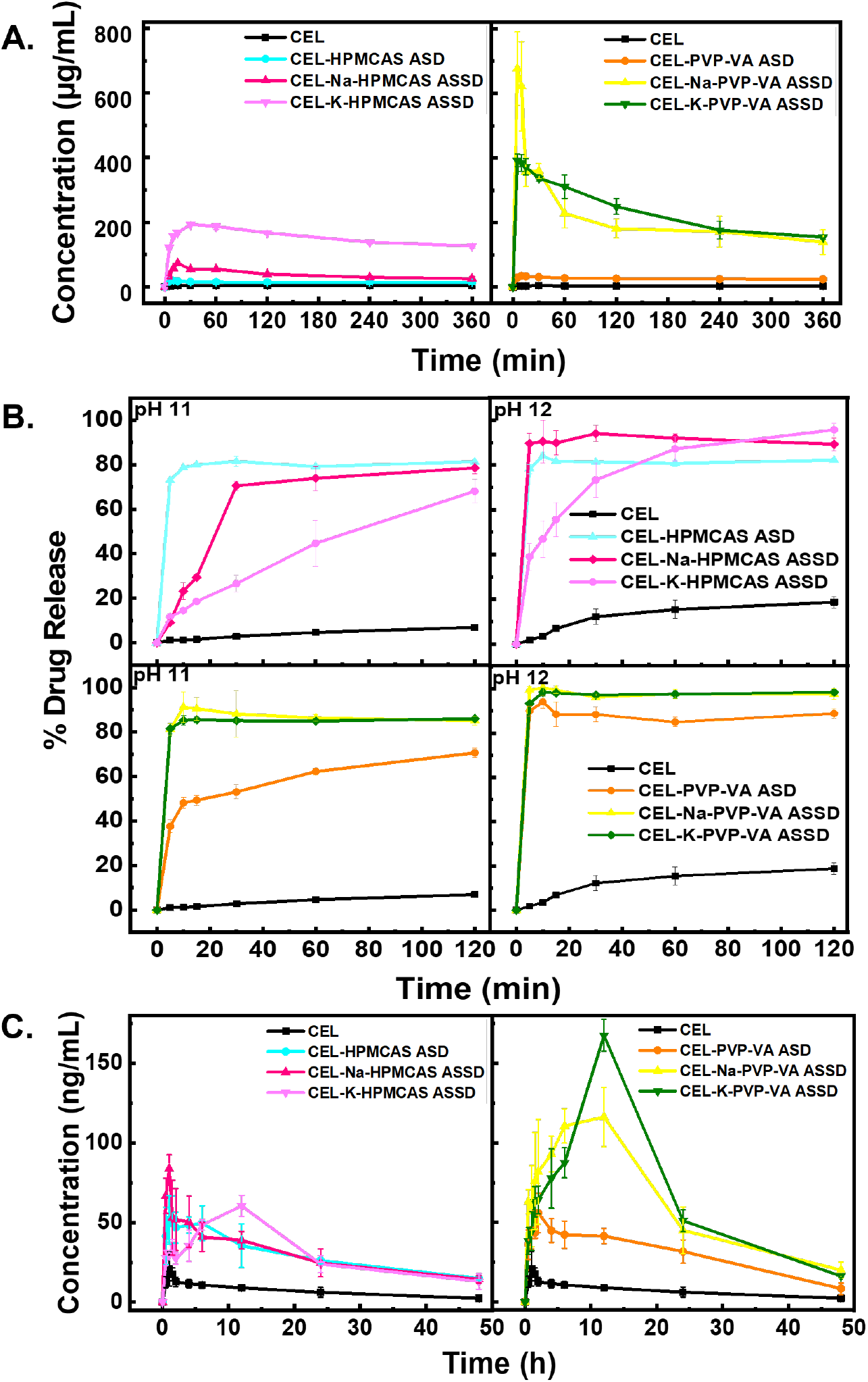
Biopharmaceutical properties of crystalline CEL and its amorphous formulations, showing the best performance for the CEL-PVP-VA salts. **(A)** Apparent solubility in aqueous medium (water) for 6 h. **(B)** Dissolution profile under sink and non-sink conditions in pH 11 and 12 buffer media, respectively, for 2 h as per OGD recommendation for dissolution performance. Measurements for (A) and (B) were made in triplicate. **(C)** Plasma concentration vs time profile from pharmacokinetic studies performed for 48 h in SD female rats (n=5; data provided in accompanying Excel file).

#### In vitro dissolution studies

*In vitro* dissolution studies were performed to understand the release behaviour of pure crystalline CEL and CEL from amorphous ASD and ASSD formulations under sink (pH 12) and non-sink conditions (pH 11). The pK_a_ of CEL is 11.1 so crystalline CEL remains unionized throughout the GIT at pH values below 11. At pH 11, CEL is ∼50% ionized and at pH 12, CEL is more ionized, causing enhanced solubility. Indeed, the equilibrium solubility of pure crystalline CEL in pH 12 dissolution medium was measured to be 569.80 μg/mL, which is much higher than the solubility at pH 11 (28.09 µg/mL). Hence, dissolution studies were performed at both pH 11 and pH 12.

##### Dissolution under sink conditions

The dissolution studies were performed under sink conditions at pH 12 using tribasic sodium phosphate buffer (1000 mL media) as recommended by the Office of Generic Drugs (OGD), USFDA for robustness and biological relevance. As per United State Pharmacopoeia (USP), the sink condition is defined as the condition in which one entire dose of the drug is solubilized in one third of the volume of the dissolution medium. Accordingly, an entire 100 mg dose of CEL should solubilize in 175 mL dissolution volume at pH 12.

Crystalline CEL showed release of 18.73 % after 120 min whereas the formulated binary CEL-PVP-VA and CEL-HPMCAS ASDs exhibited CEL release of 88.00 % and 82.17%, respectively, after 120 min. For the CEL-Na-PVP-VA ASSD, 100.00 % of CEL was released within the initial 10 min. However, the released CEL concentration declined to 97.55 % due to its de-supersaturation after 120 min. The CEL-K-PVP-VA ASSD manifested a CEL release of 98.46 %. In the CEL-Na-HPMCAS ASSD, a rapid increase in CEL release was observed in the initial 5 min followed by supersaturation maintenance over 120 min with a CEL release of 89.27 %. In contrast, the CEL-K-HPMCAS ASSD showed a gradual increase in CEL release, reaching 95.67 % after 120 min, slightly higher than for the CEL-Na-HPMCAS ASSD (Fig. 6B).The ASD and ASSD formulations followed the concept of spring and parachute effects. CEL was released rapidly above the saturation level due to its amorphous nature in the formulations (spring effect), and the supersaturation was further maintained because of the presence of polymer in close proximity to the drug, avoiding solution-induced crystallization through drug-polymer interactions (parachute effect). For the ASSD formulations, CEL solubility was enhanced due to salt formation, and the dissolution performance was better than for the corresponding binary ASD, thereby achieving improved bioavailability. The increment in dissolution rate is attributable to the dual benefit of ASSDs, amorphization and salt formation, due to the loss of crystal lattice energy and self-buffering capacity, respectively. The dissolution rate is dependent on the release of drug from the diffusional double layer around the dissolving particle in the medium. The salt alters the pH of the microenvironment around the dissolving drug particle and promotes its ionization and faster dissolution as compared to free acid or base^[53]^.

##### Dissolution under non-sink conditions

Dissolution studies under non-sink conditions were performed at pH 11 using tribasic sodium phosphate buffer. These conditions are preferred for discriminating the drug release of amorphous formulations and crystalline CEL, for direct evaluation of ASDs and ASSDs for enhancing solubility and maintaining supersaturation as supersaturation is better sustained in non-sink conditions, and for ensuring product quality and *in vitro* and *in vivo* performance^[54]^. The equilibrium solubility of CEL in this buffer was found to be 28.09 μg/mL. Therefore, the entire 100 mg dose of the CEL cannot be solubilized in one third of the volume of the dissolution medium and the system corresponds to non-sink conditions and provides a more realistic estimate of the ability of amorphous solid based formulations to increase the release of PWSD in the GIT. Crystalline CEL showed a low release rate of 6.96 % in pH 11 buffer after 120 min because of its low solubility and poor wetting behavior. Because of its hydrophobic nature, CEL tended to float on the surface of the dissolution medium during the study period. The CEL release exhibited by the CEL-PVP-VA ASD was approximately 70.62 % after 120 mins, which was considerably lower than for the PVP-VA ASSD. On the other hand, the CEL release from the CEL-HPMCAS binary ASD was higher at 81.32 % compared to that for the CEL-Na-HPMCAS and CEL-K-HPMCAS ASSDs for which the CEL release was 78.53 % and 68.08 %, respectively after 120 minutes (Fig. 6B), indicating slow and incomplete release from the HPMCAS polymeric matrix. However, both the CEL ASSD formulations displayed approximately 12-fold enhancement in the percentage of drug release compared to pure crystalline CEL.

CEL-Na-PVP-VA and CEL-K-PVP-VA ASSD formulations exhibited a similar pattern of maximum drug release, with 85.09 % and 85.98 % release after 120 mins (Fig. 6B). Moreover, a difference of approximately 40 % was observed in the release profiles of CEL-PVP-VA ASD and its ASSD for the initial 10 min, with the difference falling to 15 % after 120 min. Apart from strong ionic interactions in amorphous salts, strong CEL-polymer interactions in both the ASSDs prevented the aggregation of CEL particles from the supersaturated solution. Under non-sink conditions, the ASSD prepared with HPMCAS did not generate supersaturation greater than its binary ASD. The CEL release from both the ASSDs was gradual and the ASSD and ASD prepared using PVP-VA exhibited better drug release than HPMCAS. We attribute this difference to the higher CEL-PVP-VA miscibility (Fig S1) which results in better inhibition of CEL crystallization and prolonged maintenance of supersaturation in CEL-PVP-VA amorphous formulations (Fig S7). Moreover, due to the controlled-release and hydrophilic nature of PVP-VA, swelling of the polymer matrix occurred followed by slow erosion after complete hydration^[55–57]^, leading to more drug release.. The higher dissolution of ASSD formulations corresponded to their *in vivo* biopharmaceutical performance.

The release of CEL from CEL-Na-HPMCAS ASSD was much faster than CEL-K-HPMCAS ASSD at both pH 11 and pH 12 because of the low Tg value of the former. While the smaller Na^+^ ion might be expected to interact more strongly with CEL than the larger K^+^ ion, the K^+^ ion has a lower ionization enthalpy and may therefore have a greater tendency to form ionic interactions with CEL in the amorphous form leading to slower drug release. Whereas the CEL release profile for CEL-HPMCAS ASD is very similar at pH 11 and pH 12, for the ASSDs, the extent of CEL release is lower than the CEL-HPMCAS ASD at pH 11 and higher at pH 12, and the rate of CEL release increases from pH 11 to pH 12. This difference in CEL release behaviour might be explained by the relative differences in ionization tendencies of the salt and the drug at the two pH values, with more CEL being in the ionized form at pH 12, as well as tendency of the ASSDs to form ionic interactions with CEL in the amorphous form leading to slower drug release. HPMCAS, being rather hydrophobic in nature^[58]^, is unlikely to affect the differences in ionization of the drug and the salt at these pH values.

#### In vivo pharmacokinetic studies

The *in vivo* performance of the amorphous formulations was assessed by pharmacokinetic studies. Crystalline CEL showed a maximum plasma concentration (C_max_) of 21 ng/mL at 1 h whereas CEL-PVP-VA and CEL-HPMCAS binary ASDs showed C_max_ of 56 ng/mL (at 2 h) and 52 ng/mL (at 1 h), respectively. The CEL-K-PVP-VA and CEL-Na-PVP-VA ASSDs exhibited higher C_max_ values of 167 ng/mL and 116 ng/mL, respectively, at 12 h whereas the CEL-Na-HPMCAS ASSD attained a C_max_ of 84 ng/mL at 1 h, which further reduced with time. Also, the CEL-K-HPMCAS ASSD acquired a lower C_max_ of 60 ng/mL, even at 12 h (Fig. 6C). Thus, CEL-K-PVP-VA and CEL-Na-PVP-VA ASSDs had approximately 8.0 and 5.6-fold higher C_max_, respectively, than crystalline CEL. Moreover, the extent of enhancement in C_max_ and other pharmacokinetic parameters was much greater for the ASSD than the binary ASD, see Table 3. Both the ASSD formulations of CEL with PVP-VA possessed a higher plasma concentration of CEL for 12 h due to maintenance of supersaturation for a prolonged period (synergistic effect of amorphization and salt form), enhanced solubility and dissolution, and increased intermolecular drug-polymer interactions in the ionic state than in the neutral state, thereby inhibiting drug-drug aggregation as revealed from the experimental and *in silico* computational interaction studies. This may be ascribed to generation of CEL-rich nano-droplets or a colloidal system in equilibrium with the molecularly dissolved CEL^[59]^. Also, the PVP-VA matrix allowed higher CEL loading, and enabled carrier-controlled release for continuous dissolution of CEL through gel-like layer of PVP-VA, and absorption into the blood stream gradually over a longer time period^[57]^. However, the binary ASD showed lower supersaturation potential then the ASSD due to weak intermolecular interactions between CEL and PVP-VA, resulting in a lower C_max_ (56 ng/mL). Additionally, the formulations with HPMCAS did not prove better than PVP-VA, indicating the lower supersaturation potential of HPMCAS that is evident from its pH-dependent solubility, i.e., lower solubility at acidic pH (< 10% below pH 4 and ∼50% at or above pH 5)^[50]^. Notably, the solid dispersion of paclitaxel (PTX) with HPMCAS-MF failed to improve the oral bioavailability of PTX in Sprague Dawley (SD) rats despite maintenance of *in vitro* supersaturation^[60]^.

**Table 3.** *In vivo* pharmacokinetic parameters of crystalline CEL, and CEL ASD and ASSD formulations.

| Formulations | C<sub>max</sub><br>(ng/mL) | Fold<br>increase | T<sub>max</sub><br>(h) | AUC (ng.h/mL)<sup>#</sup> | Fold<br>increase | MRT<br>(h) | T<sub>1/2</sub><br>(h) |
| --- | --- | --- | --- | --- | --- | --- | --- |
| Crystalline<br>CEL | 20.94± 10.85 | - | 1± 0.28 | 374.09± 25.98 | - | 24.09± 2.35 | 17.5± 1.45 |
| CEL-PVP-VA<br>ASD | 55.93± 12.21 | 2.67 | 2± 0.5 | 1632.67± 226.76 | 1.79 | 23.44± 3.45 | 15.14± 3.20 |
| CEL-Na-PVP-VA ASSD | 116.23± 18.45* | 5.55 | 12± 0.0 | 3327.57± 151.68 | 8.89 | 22.56± 0.54 | 14.89± 0.36 |
| CEL-K-PVP-VA ASSD | 167.47± 10.02* | 7.99 | 12± 0.0 | 3519.76± 124.77 | 9.40 | 19.26± 0.19 | 11.12± 0.09 |
| CEL-<br>HPMCAS<br>ASD | 51.66± 15.28 | 2.46 | 1± 1.73 | 1615.63± 126.54 | 4.36 | 27.18± 3.89 | 19.11± 3.41 |
| CEL-Na-HPMCAS<br>ASSD | 83.89± 8.74 | 4.00 | 1± 0.28 | 1878.63± 101.56 | 5.02 | 34.49± 0.98 | 24.41± 0.85 |
| CEL-K-HPMCAS<br>ASSD | 60.49± 6.52 | 2.88 | 12± 0.0 | 1774.5± 192.29 | 4.74 | 26.8± 2.86 | 17.25± 2.36 |

\*p < 0.05, statistically significant difference in comparison with crystalline CEL at n=3 (n= no of animals in each group)
<sup>#</sup>The fold increase represents the fold-enhancement in the C<sub>max</sub> and AUC of prepared formulations compared to pure crystalline CEL.

The better relative bioavailability of the ASSDs compared to the ASD for PVP-VA versus HPMCAS is due to the much better performance of the PVP-VA ASSDs, which correlates with their higher solubilities and higher drug release in dissolution media. These properties are due to the more hydrophilic nature of PVP-VA, resulting in stronger interactions with of the polymer with the anionic form of CEL, as seen from the MD simulations, which is favored by the presence of salt. This higher CEL solubility and dissolution result in more drug being available in the gastrointestinal lumen for passive absorption and lead to higher bioavailability than the corresponding ASD. In contrast, the HPMCAS polymer did not provide much enhancement in CEL solubility in water and lower CEL release was observed in dissolution media. The weaker interactions of HPMCAS with the anionic form of CEL, as seen from the MD simulations, resulted in smaller differences in the solubility and dissolution profiles of the HPMCAS ASD and ASSDs, and therefore, little difference in their bioavailability.

### Evaluation of formulation stability

The physical stability of the ASSDs was evaluated at ambient conditions and compared to that of the corresponding binary ASDs. All the ASSD formulations were found to be stable for more than 11 months (352 days) at the ambient temperature and relative humidity present in the lab. Even after 11 months storage, DSC thermograms of ASSD formulations prepared with Na^+^ or K^+^ counterions exhibited T_g_ and showed no recrystallization and no melting endotherm (see Fig. S11d-g). The DSC thermograms correlated well with the optical and polarized light microscopy data which showed no birefringence in polarized mode (see Fig. S12e-h). The results support the amorphous nature of the formulations and the absence of recrystallization, indicating the prolonged physical stability of ASSD formulations. ASDs, on the other hand, showed a melting endotherm in DSC thermograms after 11 months, suggesting recrystallization occurred and indicating poor physical stability (see Fig. S11b,c). Consistently, in optical and polarized light microscopy images, birefringence was observed in the polarized mode, confirming the presence of crystallinity in the samples and the loss of the amorphous nature (see Fig. S12c,d). Hydrogen bonds in the ASDs were not able to prevent crystallization over longer time periods whereas the stronger ionic interactions in the ASSDs could. Moreover, amorphous CEL was found to be unstable 48 h after preparation at room temperature as revealed by the birefringence observed in optical microscopy (Fig S12b) and the DSC thermogram showing recrystallisation at high temperature (Fig. S4B). Thus, the ASSD formulation enhanced the physical stability of the amorphous form over long times in comparison to conventional ASDs.

## DISCUSSION

The development of next generation ASDs for enhancing the solubility of PWSDs is much needed due to problems of dose dumping and low drug loading capacity, which lead to adverse side-effects, high dosage regimens, and increased overall costs of therapy. Here, we developed advanced ASDs, namely ASSDs, of CEL, a widely used anti-inflammatory drug that is a PWSD. These ASSDs alleviate many of the problems of conventional ASDs. Although it is established that polymers and salts can play an important role in solubilizing and stabilizing PWSDs, the detailed mechanisms governing the properties of ASDs are poorly understood. The effect of polymers on intermolecular drug-drug interactions in both ionic and neutral states of the drug in aqueous phase are largely unexplored and the basis for the selection of an optimal salt-polymer combination for a particular drug is unclear. Therefore, we here performed *in silico* MD simulations of drug-salt-polymer systems in an aqueous environment and compared the results to experimental measurements of *in vitro* properties and the *in vivo* fate of the drug. Atomic detail MD simulations of such systems have only recently become feasible due to the large size and chemical heterogeneity of excipient polymers, making it necessary to simulate systems with large numbers of atoms for long times to obtain sufficient sampling. Thus, the MD simulations were carried out to explore the effects of polymers and ions on drug-drug interactions and drug aggregation, but drug crystallization processes were not simulated as they take place on much longer timescales than the nano-microsecond durations of the MD simulations. For computational feasibility, each polymer was represented by a single oligomer chain that was much shorter than the real polymer chains. While each single modeled oligomer chain had the correct relative abundance of substituents, it was not possible to capture the heterogeneity in substituent positions of the polymers used in the experiments. With advances in computing hardware and software algorithms, we expect that in the future the approach taken in this study can be expanded to more complete models with more extensive temporal, chemical and spatial sampling of the aqueous drug-salt-polymer systems, as well as consideration of further molecular components of the *in vivo* environment of the gastrointestinal tract. Nevertheless, the MD simulations described here provide a basis for probing the determinants of CEL-polymer excipient interactions. We focused on ASDs containing CEL, and one of two established polymer excipients -PVP-VA or HPMCAS – with or without Na^+^ or K^+^ salts. While classical empirical models showed adequate miscibility of CEL and both polymers, they were unable to correctly identify the best drug-salt-polymer combination. In contrast, more stable drug-polymer intermolecular interaction energies computed from the atomically detailed MD simulations were found to correlate with prolonged stability of supersaturated amorphous drug-salt-polymer systems in aqueous medium. MD simulations revealed that the PVP-VA oligomer formed a relatively more stable complex with anionic CEL than HPMCAS and experiments for ASSDs, in which the anionic form of CEL is favored, revealed that it was a better polymer than HPMCAS for inhibiting precipitation and maintaining supersaturation of CEL for a prolonged time. All simulations of CEL-containing systems were performed in the presence of NaCl and, as no direct effects of the Na^+^ ions on the CEL:CEL or CEL:oligomer interactions were observed, similar behaviour would be expected in classical MD simulations with K^+^ ions. Molecular simulations to capture the differences between these cations would likely require the use of a polarizable force field or a quantum-mechanics-based model. However, experiments did show some differences in solubility between ASSDs with the two cations, likely due to differences in the size and polarizability of the cations, whereby K^+^ has a higher polarizability than Na^+^, and their ability to polarize their surroundings, which is greater for Na^+^ than K^+[52]^. Thermal stability analysis revealed that the CEL-Na-PVP-VA and CEL-K-PVP-VA formulations exhibited higher T_g_ values, proffering good physical stability, and various *in vitro* and *in vivo* studies showed correspondingly improved biopharmaceutical performance. This novel technology of amorphous formulation of CEL facilitates *in situ* salt formation within the polymer matrix leading to high drug-loading and prolonged stability, along with ameliorated aqueous solubility, dissolution, and pharmacokinetic profiles. It thus offers the prospect of less frequent administration and lower doses with good patient compliance, and therefore fewer side effects.

It should be noted that the ASSD technology is not suitable for non-ionizable drugs due to their inability to form a salt. In addition, due to the higher hygroscopicity of ASSDs than ASDs (Fig. S14) because of their amorphous nature and the presence of the Na^+^ or K^+^ salts which are very hygroscopic^[61]^, their applicability could be challenging for more hygroscopic drugs. The ASSD technology remains to be explored for permeability enhancement of BCS class IV drugs, that have both low permeability and low solubility. Despite such limitations, the ASSD technology has potential for the formulation of PWSDs with salt-forming abilities that have compromised bioavailability.

The PVP-VA ASSDs characterized in this work may, upon further development for industrial production, find direct application for the administration of CEL due to their excellent biopharmaceutical properties and cost-effectiveness compared to formulations currently in use. Indeed, a straightforward scale-up for their production should be feasible. Although we only considered one drug and a limited number of polymer-salt combinations in this study, the combined computational and experimental approach described here is generally applicable to other classes of drugs and polymer types and can be used to aid the rational choice of drug-salt-polymer combinations for drug formulation. Moreover, we expect the physical insights gained into the determinants of amorphous formulations to be relevant for the design of drug-salt-polymer combinations for the formulation of the large class of PWSDs. Finally, the ASSD technology could pave the way for biowaivers due to the immediate release of more than 85 % of the drug from the ASSD formulations within 15 to 30 minutes of dissolution, as required by USFDA guidelines for immediate-release drug products^[62]^; this would provide the dual benefit of cost and time by circumventing the need for *in vivo* bioequivalence studies.

## MATERIALS AND METHODS

### Experimental design

The experimental and computational characterizations of the systems (drug, polymer, formulation) studied were performed in parallel and the results compared. The systems were chosen to allow systematic comparison of the effects of two polymers – HPMCAS and PVP-VA in the presence or absence of Na^+^ or K^+^ salts on the physicochemical and biopharmaceutical properties of CEL when formulated as amorphous (salt) solid dispersions. The experiments were designed with the aim of identifying the determinants of system behaviour at supersaturation in aqueous phase and of CEL bioavailability.

### Materials

CEL form III was provided as a gift sample by Jubilant Generics Ltd. (Noida, Uttar Pradesh, India). PVP-VA 64, manufactured by BASF (Germany) and HPMCAS-MF grade, manufactured by Shin Estu (Japan) was generously provided by Signet Chemical Corporation Pvt. Ltd. (Mumbai, India). NaOH and KOH were procured from Himedia Laboratories Pvt. Ltd. (Mumbai, India). All other chemicals used in the experiments were of analytical grade and used as such.

### Determination of drug-polymer miscibility

Estimation of δ by the Fedors and HVK methods is the conventional approach to calculate drug-polymer miscibility in order to prepare thermodynamically stable amorphous solid based formulations^[63, 64]^. The drug and polymer are found to be miscible if both have similar δ values with the difference in the solubility parameters (Δδ) being less than 7 Mpa^1/2^. The Hildebrand solubility parameter for the CEL-PVP-VA and CEL-HPMCAS systems was calculated using Fedors method from the cohesive energy density (CED) using Equation 1, where E_v_ is the energy of vaporization, V_m_ is the molar volume, and CED is the cohesive energy per unit volume^[60]^. The HVK method considers the dispersive forces, interactions between polar groups, and hydrogen bonding groups as shown in Equation 2, where δ_d_, δ_p_ and, δ_h_ are the dispersive, polar, and hydrogen bonding solubility parameter components, respectively. These components can be further calculated using Equation 3, where F_di_ (dispersion component), F_pi_ (polar component) and E_hi_ (hydrogen bonding energy) are the group contributions for different components of structural groups and V is the group contribution to molar volume^[63]^. Further, the lattice-based F-H interaction parameter, χ was estimated using the δ value describing the change in Gibbs free energy before and after mixing of drug and polymer as shown in Equation 4, where χ refers to the square of the difference in δ value of the drug and the polymer, calculated from group contribution methods at 25 °C, R is the gas constant, T is the absolute temperature, and V_site_ is the volume per lattice site^[63]^.

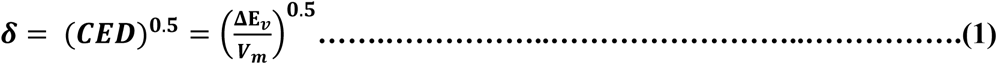

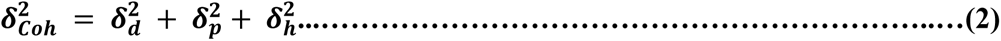

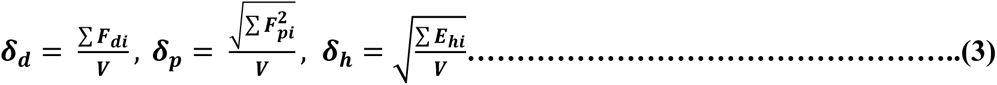

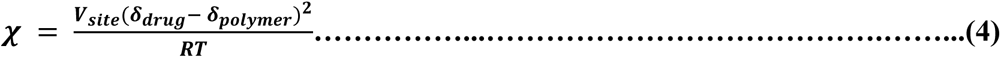

Experimental determination of drug-polymer miscibility was through melting point depression studies where PMs were prepared at different drug:polymer ratios of 100:0, 95:05, 90:10, 85:15, 80:20, 75:25 and 70:30 by geometric mixing with a spatula in a mortar followed by heating at the rate of 10 °C/min in DSC and the depression in the melting point of the PM was observed.

### Generation of ASSD and binary ASD formulations

The generation of ASSD formulations was previously explained in detail^[27]^. Briefly, CEL (pKa 11.1) and counterions {K^+^ (KOH (pKa 14.7)) or Na^+^ (NaOH (pKa 15.7))} were dissolved in methanol in a 1:1 molar ratio to generate Na^+^ and K^+^ salt solutions. The salt formation is attributable to the pK_a_ difference of more than 2 between CEL and the counterions. The two polymers, HPMCAS and PVP-VA, were dissolved separately in another part of methanol. The salt-containing solutions were then separately added to each polymer-containing solution in order to obtain a drug-polymer mass ratio of 6:4 and mixed well by magnetic stirring (see Fig. S2). The resulting solutions were then subjected to drying using a laboratory scale spray dryer (Buchi B-290, mini spray dryer, Buchi Labortechnik AG, (Switzerland)) set at the parameters of inlet temperature 80 °C, outlet temperature 50 °C, feed pump 12%, aspiration 85%, and air atomization pressure 400 L/h. Binary ASDs were also generated using the same set of parameters and technique by mixing CEL and HPMCAS or PVP-VA in a 6:4 mass ratio in methanol. Residual solvent was removed from both ASDs and ASSDs by storing in a vacuum oven at 30-35 °C for 5 to 6 h.

### Characterization methods

The concentration of CEL for solubility and dissolution studies was analysed using a validated high-performance liquid chromatography (HPLC) method employing an HPLC system (Shimadzu Corporation, Kyoto, Japan) with a PDA detector (SPD-M10AVP) and LC solution software as described previously^[27]^. Optical imaging of powder samples was carried out to determine the morphology using a polarized light microscope (Leica Microsystems GmbH, (Wetzlar Germany)), and photomicrographs were acquired in both optical and polarized modes at 20x magnification using Leica IM 50 (version 1.20) software to determine the crystalline or amorphous nature of samples. The nature of the samples was also confirmed by DSC and PXRD. The sample preparation for DSC studies was done as described previously^[27]^. A Q2000 DSC (TA Instruments, New Castle, Delaware, USA) with an attached refrigerated cooling system and operating with Universal Analysis software version 4.5A (TA Instruments, (New Castle, Delaware, USA)) was used. Dry nitrogen at a 50 mL/min flow rate was used to purge the sample cell. The instrument was calibrated for temperature and heat flow using a high purity indium standard. Accurately weighed samples (4-5 mg) in aluminium pans were analysed by heating at a rate of 20 °C/min from 25 °C to 200 °C. The T_g_ and endothermic transitions were reported as the midpoint temperatures. Powder X-ray diffractograms for crystalline CEL and the generated ASSDs and ASDs were acquired at ambient conditions employing a Rigaku Ultima IV fully automatic high resolution X-ray diffractometer system (Rigaku Corporation, (Tokyo, Japan)) equipped with an X-ray radiation source Cu Kα tube at 1.54 Å wavelength that was operated at 40 kV/40mA, with a divergent slit of 0.5°, a divergent height limit slit of 10 mm, a scattering slit (SS) of 8.0 mm, and a receiving slit (RS) of 13 mm. All the samples were scanned continuously at a step size of 0.02° and scan rate of 15 seconds, over the range from 3° to 50° at an angle 2θ.

The molecular level drug-polymer interactions in the formulations were investigated using high-end analytical techniques. FTIR spectra were recorded on an FTIR multiscope spectrophotometer (PerkinElmer, (Buckinghamshire, UK)) equipped with spectrum v3.02 software by a conventional KBr pellet method. The Micro-Raman Jobin Yvon HORIBA LABRAM HR-800 visible system (Kyoto, Japan) equipped with a 473 nm diode laser source, 1800 lines/mm grating, and an Olympus BX41 microscope was used to perform Micro-Raman spectroscopy. Raman spectra were recorded in back-scattering mode at room temperature with a pinhole size of 50 μm, slit width of 100 μm and exposure time of 10 seconds. Proton NMR was carried out for all the samples dissolved in deuterated DMSO-d6 using a Bruker 500 UltraShield NMR spectrometer (Billerica, Massachusetts, USA) functioning at 500 MHz.

Supersaturation and precipitation studies were conducted with a UV spectrophotometer (SpectraMax® M2 series) and software SoftMax Pro 6.2.1 (Molecular Devices, United States). The sample was immersed in 25 mL of water (aqueous media) at 37 °C for 4 hours by cuvette method. Plain crystalline CEL previously dissolved in methanol at 5 mg/mL concentration and drug-polymer samples (ASSD and binary ASD) containing the equivalent of 5 mg/mL CEL were added to the aqueous media while stirring with a magnetic stirrer and the samples were analysed at the absorbing wavelength of 252 nm at reading intervals of 30 seconds.

### Molecular modelling

The drug-polymer interplay at the molecular level was elucidated by MD simulation to address the role and specificity of the polymer in preventing drug-drug aggregation in supersaturated systems. MD simulations were performed for both neutral and anionic CEL in the presence and absence of polymers, estimating the interaction energies between oligomer:CEL and CEL:CEL complexes. The molecular models of CEL, PVP-VA and HPMCAS were generated for further simulation studies. The coordinates of the neutral form of CEL were retrieved from the crystal structure in the Cambridge Crystallographic Data Centre (CCDC)^[46, 65]^. To model the anionic state of CEL, a hydrogen atom was removed from the sulfonamide group. A PVP-VA 34-mer (Fig. 1B) was generated using Materials Science (Schrödinger version 2020) by randomly assembling 18 units of vinylpyrrolidone (VP) and 16 units vinyl acetate (VA). The mass-ratio of these two units (VP:VA) in the model polymer was roughly equivalent to 6:4. A model of a hypromellose (HPMC) 20-mer was kindly provided by Prof. Ronald G. Larson, (Department of Chemical Engineering, University of Michigan, Ann Arbor, Michigan 48109-2136, USA). We removed 5 monomeric units and then converted the side chain hydroxyl (-OH) group of HPMC to acetyl, deprotonated succinoyl, methoxy and hydroxypropoxy groups using Maestro (Schrödinger version 2020) to build a HPMCAS 15-mer. We chose the side-chain substitutions corresponding to the composition of M-grade HPMCAS^[66]^ (Fig. 1C). Consequently, the modelled 15-mer-M-grade HPMCAS polymer contained 45 substituents having 5 different functional groups in the following proportions (given as % of the number of functional groups and by weight (%wt)): methoxy (64.4%; 25%wt), hydroxypropoxy (8.9%; 8.3%wt), acetyl (11.1%; 8.3%wt), succinate (8.9%; 13%wt) and hydroxyl groups or no substitution (6.7%; 1.3%wt).

### Molecular parameterization

GAFF^[67]^ parameters were assigned to CEL and the polymers. Partial atomic charges for both CEL and the polymers were generated by the RESP^[68]^ method as implemented in Antechamber in the AMBER version 14 software package^[69]^ with the Gaussian 09^[70]^ software with B3LYP-6-31G*/HF-6-31G* level of theory^[71–74]^ being used for the electrostatic potential (ESP) calculations. For the polymers, geometry optimization followed by ESP calculations for the entire oligomers would be too computationally demanding. Therefore, RESP charges were computed for each monomeric unit separately. For this purpose, the monomeric units were built with blocking groups to account for the adjacent units. For PVP-VA, six non-terminal and two terminal (see Fig. S10) monomeric units were modelled. Likewise, for HPMCAS (Fig. 1C), 13 non-terminal monomeric units were generated by adding an isopropyl group to each of the two terminal oxygen atoms, and 2 terminal monomeric units were generated by adding an isopropyl group to the first and last terminal oxygen atom of HPMCAS. To maintain charge neutrality of the oligomers, the residual charge (-1.646e for the PVP-VA 34-mer and +1.926e for the HPMCAS 15-mer) after assigning the RESP charges to the monomers, which is due to the capping of the terminal atoms of the monomers, was distributed equally onto all the atoms in the oligomers. The TIP3P^[75]^ model was used for water molecules.

### Refinement of polymer models by MD simulation

To structurally refine the polymer models, all-atom MD simulations in explicit solvent were carried out for the two representative model structures of PVP-VA and HPMCAS. The polymers were immersed in cubic periodic boxes using the Gromacs^[76]^ insert-molecules command and then TIP3P water molecules were added to the system using the tleap module of AMBER. To avoid periodicity artifacts, the box dimensions were greater than the lengths of oligomers (see Table 1). 4 excess Na+ ions were added to neutralize the HPMCAS oligomer. The MD simulation protocol is given below. The last snapshot from each trajectory was extracted and used as the starting structure for drug-polymer simulations. The simulation lengths, box sizes and the numbers of water, ions are listed in Table 2 (systems a and b).

### Simulation of CEL aggregation

To simulate a supersaturated solution of CEL, we generated periodic boxes with the same number of CEL molecules as used for the drug-oligomer systems described below. 11 and 13 CEL molecules, which were modelled in either neutral or anionic forms, were inserted randomly in cubic periodic boxes of water molecules with a box-length of 106 and 112 Å, respectively, using the Gromacs insert-molecules and gmx solvate commands. Na^+^ ions were added to neutralize each system and thus the Na^+^:CEL mass-ratio for each simulated system with anionic CEL molecules was 0.0603:1, while in the experiments, the NaOH:CEL ratio was 0.105:1. Note that the hydroxide anions (OH^-^) are expected to extract the proton from the CEL sulfonamide nitrogen at higher pH values, and therefore the simulated systems did not contain hydroxide ions. In addition, further Na^+^ and Cl^-^ ions were added to all the simulation boxes to generate ∼0.01M ionic solutions (see Table 2). Then, the system was set up for simulation with AMBER using the tleap program for which it was necessary to first remove the hydrogen atoms from the water molecules and change the residue name of water from “SOL” to “WAT”. All-atom explicit solvent energy minimization and MD simulations were carried out for all the systems and the aggregation of CEL was monitored. The simulation lengths, box sizes and the numbers of water, ions are given in Table 2 (systems c-f).

### Simulation of CEL-polymer systems

For each of the CEL-oligomer combinations listed in Table 2 (systems g-j), one oligomer and several CEL molecules were placed randomly in a periodic simulation box using the Gromacs insert-molecules command, followed by packing of the boxes with water molecules and ions using the gmx solvate command.

The amounts of the constituents of the periodic simulation boxes were chosen to maintain the experimental CEL:polymer ratio and to mimic the relevant conditions *in vivo* as much as possible while minimizing the number of particles simulated for computational feasibility. As described above, we modelled a single chain of each polymer of sufficient length to contain its defining substituents in the ratios used in the experiments. We then computed the oligomer’s molecular weight and the number of CEL molecules approximately corresponding to the experimental drug-polymer mass ratio of 6:4 (1:0.67) in the ASDs and ASSDs. The molecular weights of the model PVP-VA 34-mer, M-grade HPMCAS 15-mer, and CEL are 3375.5, 3691.5, and 381.3 Da, respectively. Hence, to mimic the experiments, the CEL:oligomer mass ratio in our model systems was chosen to be 1:0.68 for PVP-VA and 1:0.88 for HPMCAS (the higher relative amount of HPMCAS was used because the 6:4 ratio for the experiments had not been finalized at the time the simulations were set up, but gives similar amorphous behavior, see e.g. Fig S5B for a 5:5 ratio). For assigning the solvent box size, we considered the concentration of a dose of CEL in the stomach. A typical dose of CEL is 100 to 200 mg given BD (twice a day) with or without food (https://go.drugbank.com/drugs/DB00482). We considered the highest single oral dose, 200 mg CEL, and the lowest water content: the 35 ml of an empty stomach ^[77]^.Thus, the stomach is a supersaturated system with CEL concentration 1650 times higher than its experimental equilibrium solubility (3.46 mg/L)^[27]^. Even considering half the dose of CEL and administration of the drug with water, the CEL concentration would still be supersaturated in the stomach. In order to restrict the simulation box size, we approximated the maximal CEL concentration. Thus, one PVP-VA 34-mer and 13 CEL molecules were inserted in a cubic box of length 112 Å, and one HPMCAS 15-mer and 11 CEL molecules were inserted in a cubic box of length 106 Å. Afterwards, Na^+^ and Cl^-^ ions were added (see Table 2), and the systems were re-solvated with the tleap module of AMBER as aforementioned and then, all atom explicit solvent energy minimization followed by MD simulations and post-facto trajectory analysis were performed.

### MD simulation protocols

All the systems were energy minimized using the AMBER v14 and v20 software^[68]^. Energy minimization was carried out by restraining non-hydrogen solute atoms with a force constant that gradually decreased from 1000 to 0 kcal/mol·Å^2^. During energy minimization, the maximum number of cycles was set to 14000 steps: 1400 steps steepest descent followed by 12600 steps conjugate gradient. Minimization was stopped when the root mean square energy gradient for the input coordinates was less than 0.00001 kcal/mol/Å.

MD equilibration and production runs were performed using the AMBER v20 software starting with the energy minimized coordinates. Each system was first equilibrated at constant volume and temperature (NVT) (with the AMBER parameter NTB=1) for a total of 16 ns with a time step of 2 fs at 300 K using the Particle Mesh Ewald (PME) method for long range electrostatic interactions with a non-bonded cut-off of 10 Å. The SHAKE algorithm^[78]^ was imposed to constrain all the bonds to hydrogen atoms. The Langevin dynamics method with a damping coefficient of 5/ps was used for temperature control. The first 6 ns of equilibration was carried out by gradually reducing the restraint force constant from 100 to 10 kcal/mol·Å^2^ on all non-hydrogen solute atoms. Then the remaining 10 ns of equilibration were run without harmonic restraints. The subsequent production runs were performed for times depending on the convergence of structural parameters, with a time step of 2 fs and a temperature of 300 K in an NPT ensemble (constant number of particles, pressure and temperature were maintained). All simulations were run in duplicate or triplicate with the same initial coordinates but with different velocities assigned. Coordinates were written at 2.5 ps intervals throughout the simulations.

### Analysis of MD trajectories

Analysis of the trajectories was performed using AMBER CPPTRAJ^[79]^. RMSD values of the system components were computed for the frames collected along the simulations. The radius of gyration of the oligomer was computed and the hydrogen bond and radial distribution function (RDF) analysis carried out for the equilibrated part of each trajectory (see Table S1). Hydrogen-bond occupancy was defined from the proportion of frames analysed at which the distance between the respective donor and acceptor atoms was ≤ 3.2 Å and the donor-H…Acceptor angle was ≥ 158°.

MM/GBSA solvation and interaction free energy^[80–82]^ calculations were carried out for frames extracted at 100 ps intervals from the equilibrated parts of the trajectories (Table S1). To compute the drug-drug aggregation energy using the MM/GBSA method, for each extracted frame, we considered each CEL molecule, one at a time, in the drug aggregate (i^th^ celecoxib where “i” is an integer number) and then computed the interaction energy between this molecule and the rest of the aggregate. CEL molecules that did not form parts of aggregates were neglected. The CEL-CEL interaction energy was the value computed by averaging over all the CEL molecules in aggregates in the simulated system and over all the extracted frames. The CEL-oligomer interaction energy was estimated similarly by, for each frame, computing the interaction energy of each CEL molecule with the rest of the complex, i.e.; oligomer and all the remaining CEL molecules, and then subtracting the CEL:CEL aggregation energy from the energy of the CEL:oligomer-CEL system to give the CEL:oligomer interaction energy. The final CEL-oligomer interaction energy was then obtained as the average interaction energy over all frames and overall CEL molecules in aggregates for each replica trajectory. The average and standard deviation of the MM/GBSA interaction free energy along the replicas were then computed and checked by bootstrapping (see Table S3).

Xmgrace (plasma-gate.weizmann.ac.il/Grace/) was used for plotting. Molecular visualization was performed with PyMOL (https://pymol.org/2/) or VMD^[83]^.

### Equilibrium solubility, apparent solubility and *in vitro* dissolution studies

The equilibrium solubility of CEL at pH 11 and pH 12 was measured by performing solubility studies in a shaker water bath at 37 ± 0.5°C and 100 RPM. An excess amount of CEL was added in 10 mL volume of buffer and kept in the shaker water bath for 72 h to attain complete equilibrium. Then, 1 ml of sample was withdrawn and centrifuged at 15,000 RPM for 15 min. Supernatant was taken, diluted appropriately and the concentration of CEL was determined by HPLC. Apparent solubility studies were carried out in distilled water, maintained at 37 ± 0.5 °C using a shaking water bath operated at 100 rpm. An equal amount of CEL and amorphous formulations equivalent to 20 mg CEL were added to 15 mL of water separately and aliquots of 1 mL were withdrawn at predetermined time points up to 6 h. All the samples were immediately centrifuged at 15,000 rpm for 10 min and the supernatant was collected for analysis. *In vitro* dissolution studies were conducted for the prepared ASSD formulations, the binary ASDs, the PM [of CEL, polymers and NaOH/KOH mixed geometrically in a mortar using a spatula], and crystalline CEL using a tablet dissolution apparatus (USP 37, type II) (TDT-08L, Electrolab, (Mumbai, India)) with an autosampler. The dissolution medium used was 0.04 M tribasic sodium phosphate at pH 12 (under sink conditions) [recommended by the OGD, USFDA] and at pH 11 (under non-sink conditions) without surfactant (SLS). A dose equivalent to 100 mg of CEL was added to 1000 ml of dissolution medium maintained at 37 ± 0.5 °C and at 50 rpm. A sample volume of 5 mL was withdrawn at predetermined time intervals up to 120 min, filtered through a 0.22 μm syringe filter (Nylon, Mdi membrane filters, Advanced Microdevices Pvt. Ltd., (Ambala Cantt, Punjab, India)), and the same volume of dissolution medium was supplied instantly after sample withdrawal to maintain sink conditions, whereas no replacement of media was done for non-sink conditions. The CEL concentration in the collected samples after suitable dilution for both solubility and dissolution studies was determined by the validated HPLC method detailed in Tables S4 and S5 and Fig. S13.

### *In vivo* pharmacokinetic studies

*In vivo* pharmacokinetic studies were conducted in SD rats (female, weight: 200 ± 30 g). The CPCSEA (Committee for the Purpose of Control and Supervision of Experiments on Animals) guidelines were followed for performing animal studies. The protocol for animal studies (IAEC/19/51-R) was sanctioned by the Institutional Animal Ethics Committee, NIPER, Mohali, India. All the animals were exposed to 12 h light-dark cycles at 25 °C and 60 % RH for one week and were fasted for 12 h with free access to water before the start of experiments. 7 groups, each consisting of 5 animals, were used for testing seven formulations viz. CEL-Na-PVP-VA and CEL-K-PVP-VA; ASSD, CEL-Na-HPMCAS and CEL-K-HPMCAS; ASSD, binary CEL-PVP-VA and CEL-HPMCAS; ASD, and pure crystalline CEL. All formulations were administered orally via oral gavage at a dose of 10 mg equivalent of CEL/kg of rat body mass. Blood samples of approximately 150-200 μL were collected from the tail vein at predetermined time points up to 48 h. Plasma was separated from the blood samples by centrifugation at 10,000 rpm for 10 min at 4 °C and stored at −40 °C before analysis. The pharmacokinetic parameters, C_max_, T_max_, mean residence time (MRT) and t_1/2_, were derived from the mean plasma concentration−time profiles using the Thermo Kinetica software (V5.0, Thermo Fischer Scientific, USA).

### Stability studies

ASSDs and binary ASDs were evaluated for physical stability at ambient conditions in closed containers after a period of 11 months. Samples were evaluated by polarized light microscopy for detection of birefringence, and by DSC for detecting melting endotherm, indicating the presence of crystallinity in amorphous formulations.

### Statistical analysis

The statistical data analysis was done by one-way analysis of variance (ANOVA) using GraphPad Prism 9 software, version 9.1.0 (221) (GraphPad Software, LLC. San Diego, USA). The results were considered statistically significant when p < 0.05.

## Supporting information

Data for Fig 6

Datat for Fig S5

## DATA AND MATERIALS AVAILABILITY

All data are available in the main text or the supplementary materials or the repository at Zenodo at: https://doi.org/10.5281/zenodo.8228709. Materials are available from the authors on reasonable request.

## CODE AVAILABILITY

All scripts for generation and analysis of the molecular simulations are available in the repository on Zenodo at: https://doi.org/10.5281/zenodo.8228709.

## Acknowledgments

We thank Prajwal P. Nandekar for his contributions in the initial phases of this study, Giulia Paiardi (HITS) for helpful discussions and assistance with data archiving, Stefan Richter (HITS) for technical support for the computational work, and Outi Salo-Ahen (Abö Akademi University, Turku, Finland) for helpful comments on the manuscript.

## Funding

German Federal Ministry for Education and Research (BMBF) 01DQ19004 (RCW) Klaus Tschira Foundation (GM, NS, CD, RCW)

Baden-Württemberg bwHPC for computing resources (RCW)

German Research Foundation (DFG) grant INST 35/1134-1 FUGG for computing resources (RCW)

Department of Biotechnology, New Delhi, BT/IN/BMBF-BioHR/34/ATS/2018-19 (ATS).

## Author contributions

Conceptualization: SM, GM, ATS, RCW

Experimental Investigation: SM, RS, PJ

Computational Investigation: GM, NS, CD

Experimental Data Analysis: SM, RS, ATS

Computational Data Analysis: GM, NS, CD, RCW

Visualization: SM, RS, GM

Supervision: ATS, RCW

Writing—original draft, review and editing: RS, GM, SM, ATS, RCW

