## Supplementary Material for "Comparative analysis of drug-salt-polymer interactions by experiment and molecular simulation improves biopharmaceutical performance"

**This PDF file includes:**

Figs. S1 to S14
Tables S1 to S5

**Other Supplementary Materials for this manuscript include the following:**

Excel file of pharmacokinetic data for Fig 6C and Table 3.
Excel file of powder X-ray diffractogram Data for Fig. S5A.
Input, parameter, coordinate and analysis files for all computations have been deposited at zenodo: https://doi.org/10.5281/zenodo.8228709

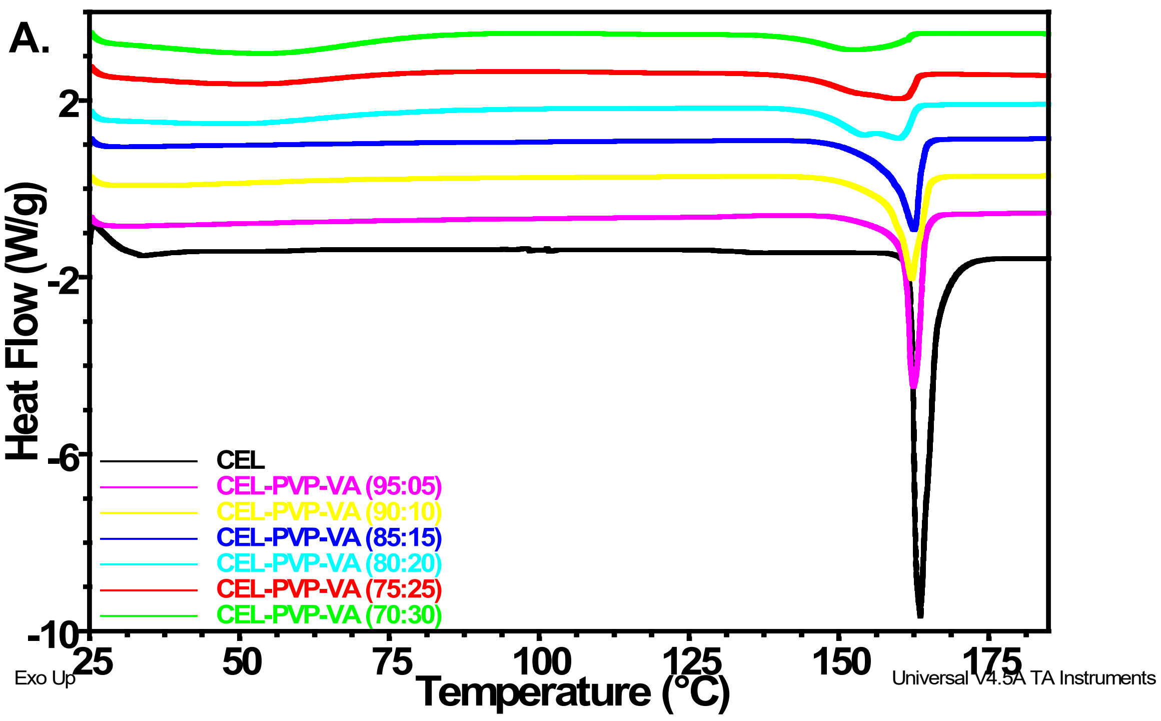


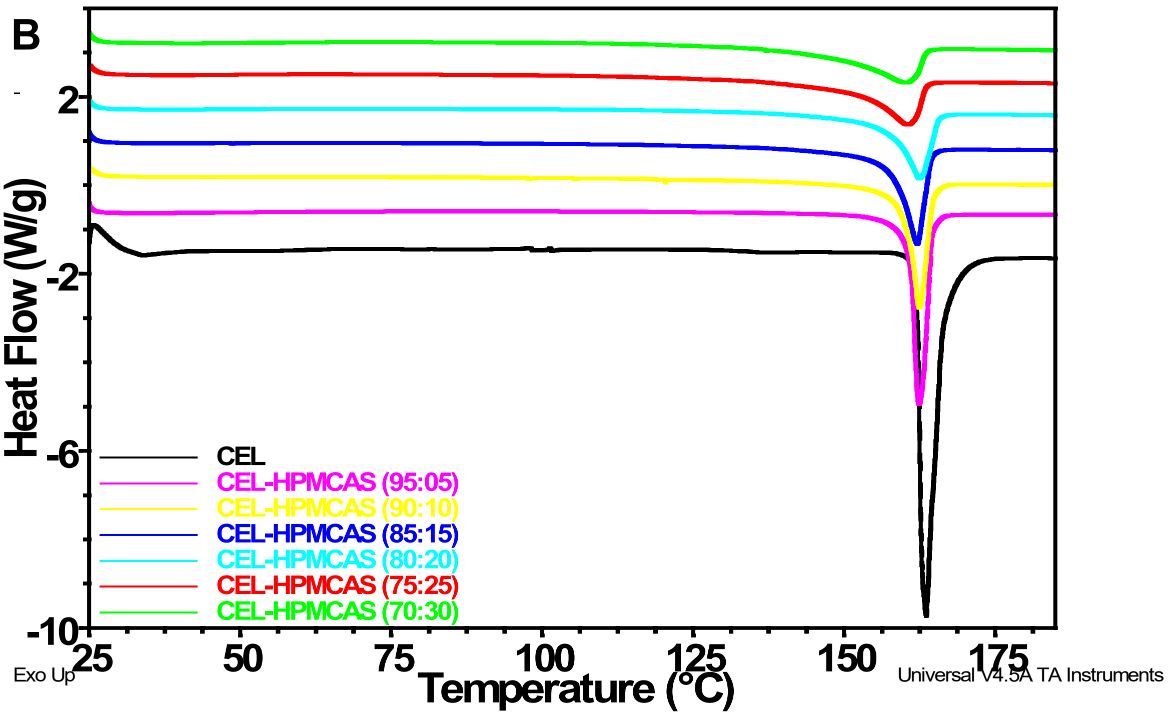


**Fig. S1.**

DSC thermograms of physical mixtures (PM) of (A) CEL-PVP-VA, (B) CEL-HPMCAS showing more depression of the melting point with PVP-VA than HPMCAS, indicating that PVP-VA has higher miscibility with CEL than HPMCAS, and can therefore form more stabilized ASDs (Fig S11).

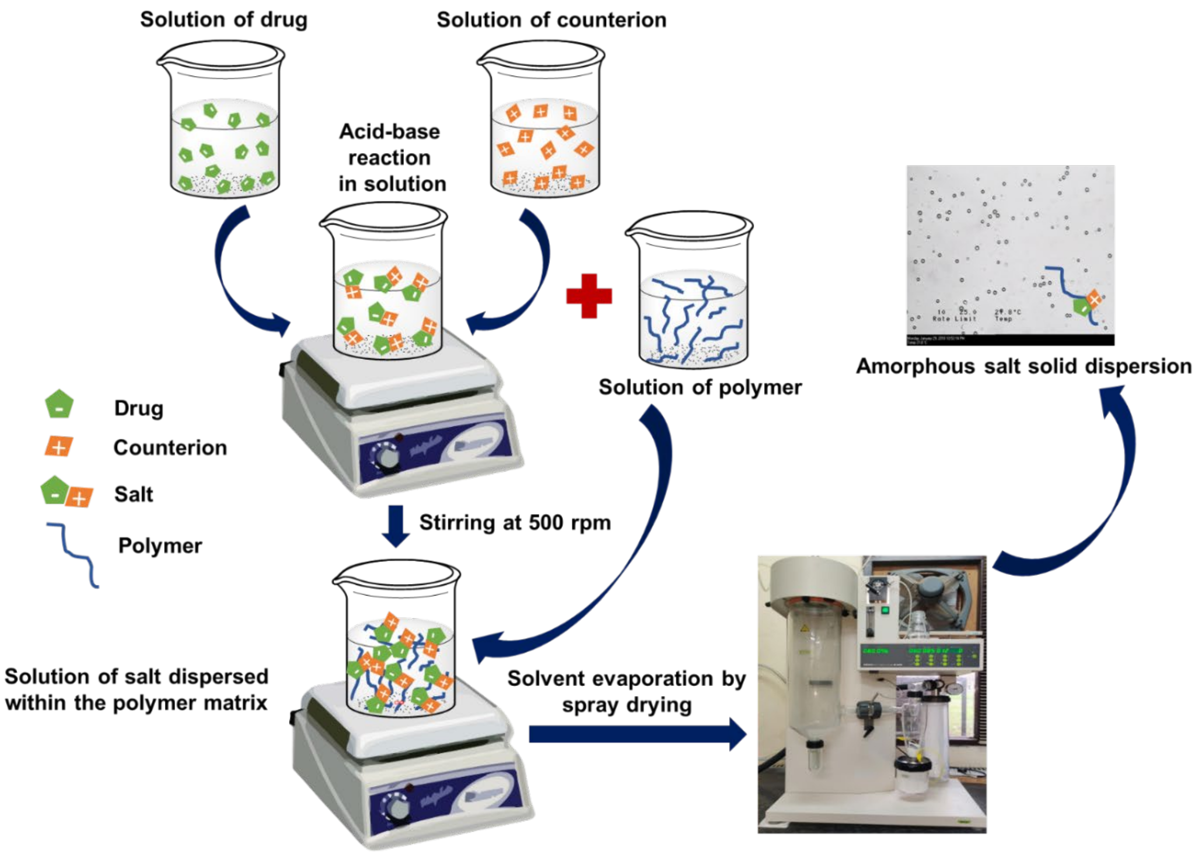


**Fig. S2.**

Schematic representation of the preparation of an *in situ* salt of a drug within a polymer matrix using a magnetic stirrer and a spray dryer.

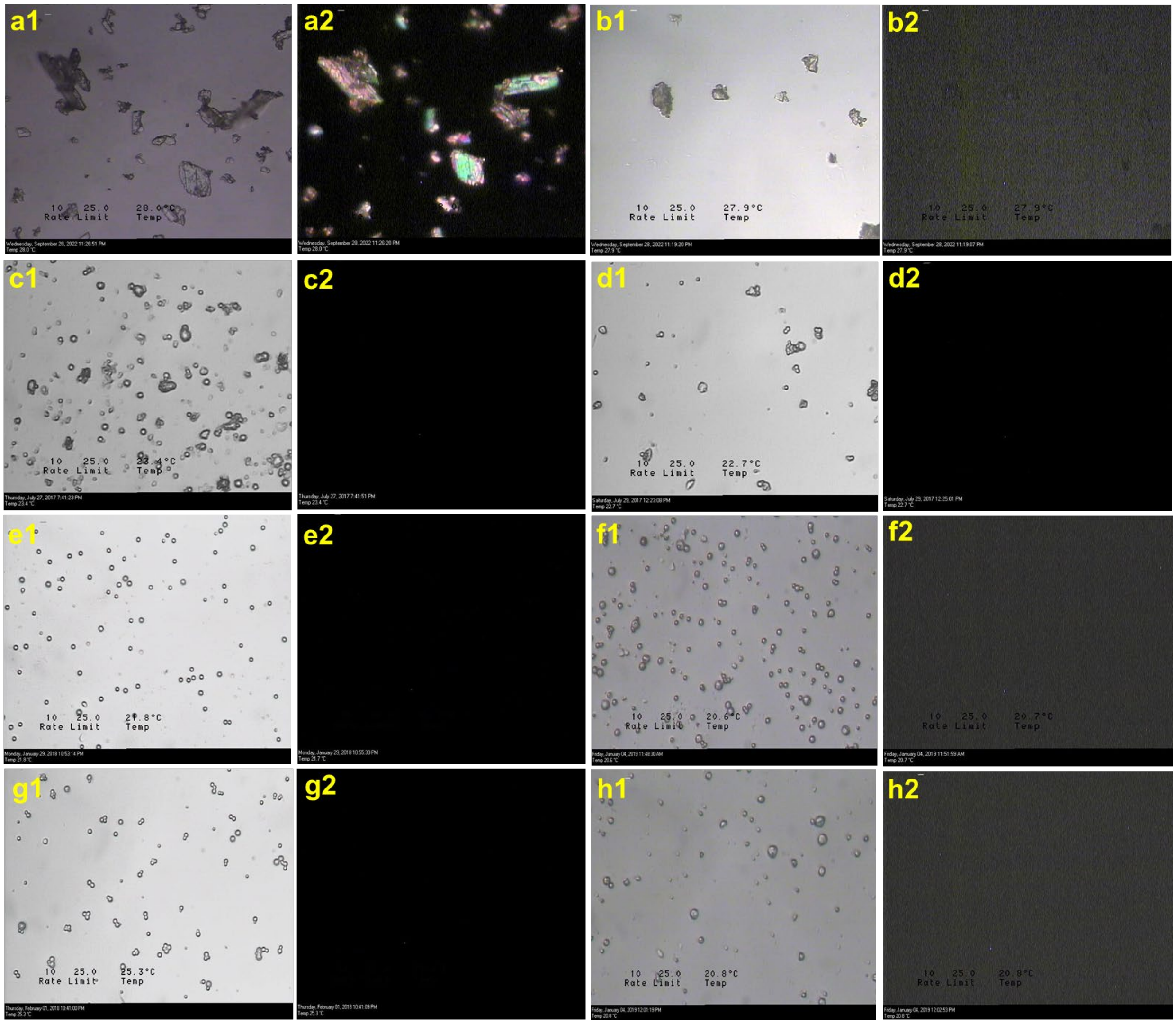


**Fig. S3.**

Optical microscopy of (a) crystalline celecoxib (CEL), (b) amorphous (AMR) CEL, (c) CEL-PVP-VA ASD, (d) CEL-HPMCAS ASD, (e) CEL-Na-PVP-VA ASSD, (f) CEL-K-PVP-VA ASSD, (g) CEL-Na-HPMCAS ASSD, and (h) CEL-K-HPMCAS ASSD. For each case, images are shown for non-polarised (1) and polarised (2) modes. The latter show birefringence in crystalline CEL (a) but no birefringence in amorphous CEL and the CEL formulations (b-h).

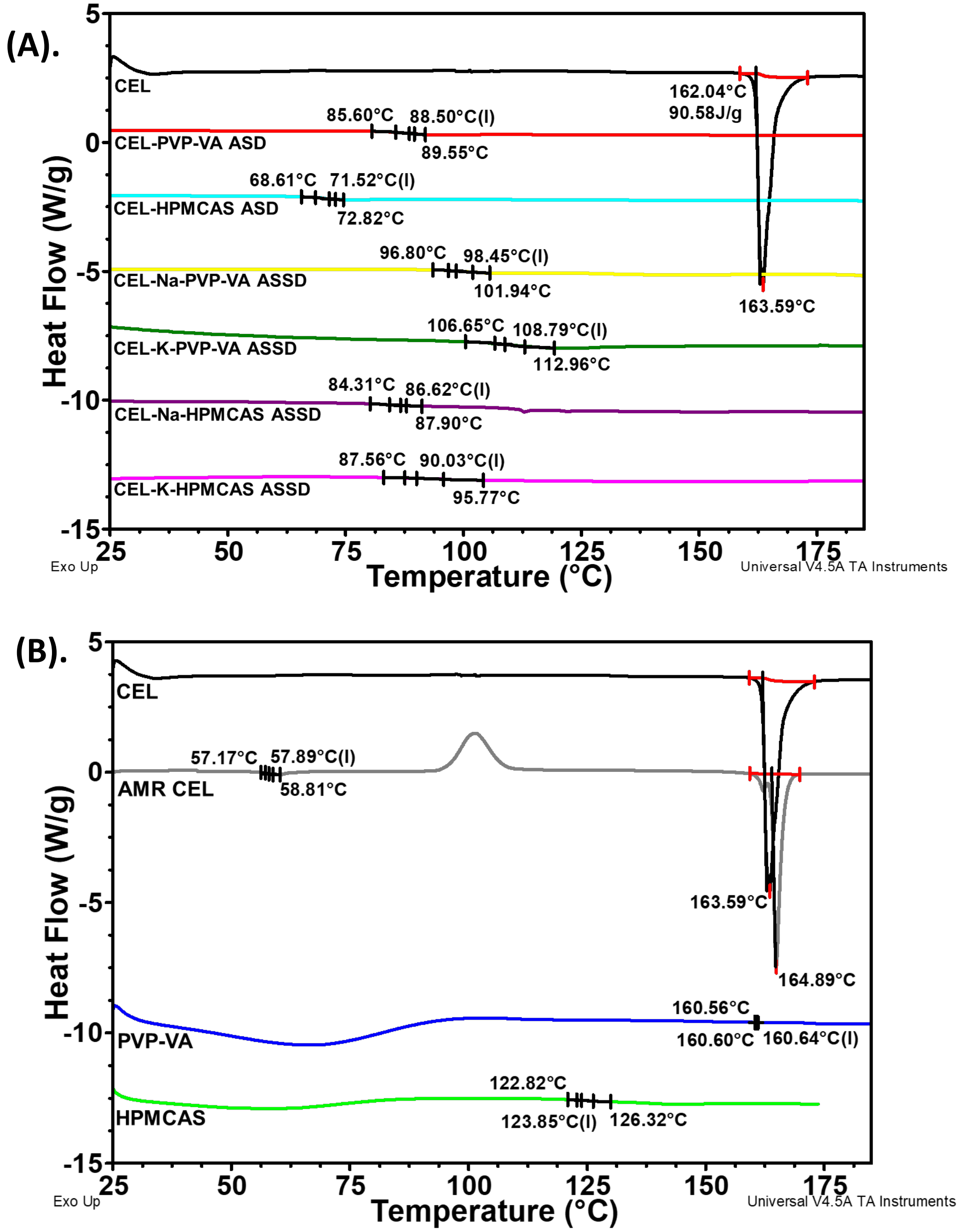


**Fig. S4.**

DSC thermograms of CEL, polymers and amorphous CEL-polymer formulations. (A) crystalline CEL, CEL-PVP-VA and CEL-HPMCAS ASDs, CEL-Na-PVP-VA, CEL-K-PVP-VA, CEL-Na-HPMCAS and CEL-K-HPMCAS ASSDs. The thermograms show a single sharp melting peak for crystalline CEL at 163.59 °C and higher glass transition temperature ($T_g$) values for amorphous formulations containing counterions than conventional ASDs, with CEL-K-PVP-VA ASSD having the highest $T_g$, which is attributed to strong intermolecular interactions. (B). Amorphous (AMR) CEL showing a low $T_g$ (57.9 °C) and recrystallisation at higher temperature, and the PVP-VA and HPMCAS polymers having higher $T_g$ values. (I) denotes the integrated value of $T_g$, which is the temperature at the mid-point of the heat-flow transition.

(A).

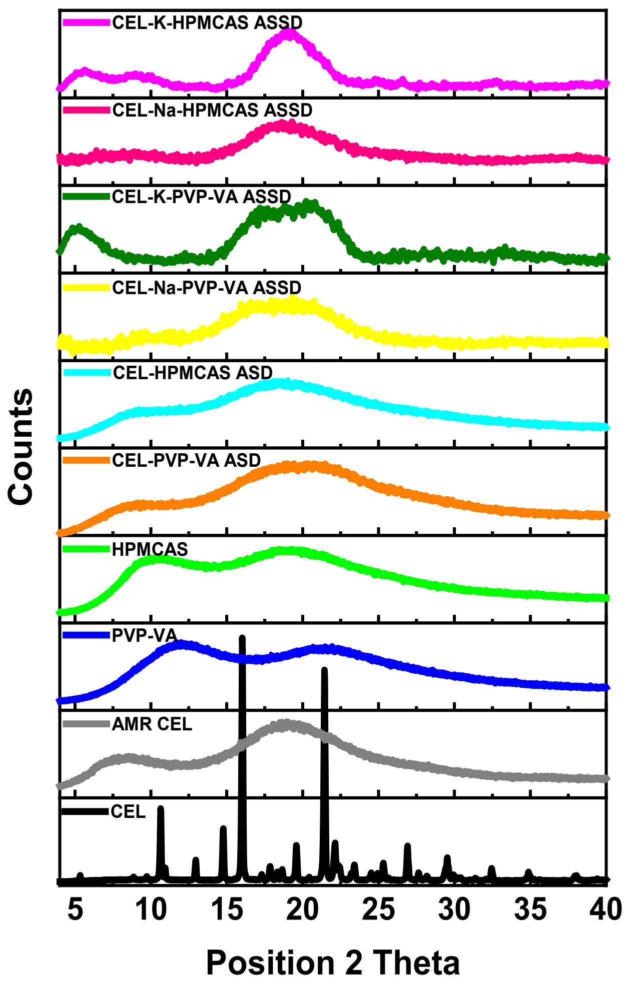
CEL-K-HPMCAS ASSD
CEL-Na-HPMCAS ASSD
CEL-K-PVP-VA ASSD
CEL-Na-PVP-VA ASSD
CEL-HPMCAS ASD
CEL-PVP-VA ASD
HPMCAS
PVP-VA
AMR CEL
CEL
Counts
5
10
15
20
25
30
35
40
Position 2 Theta

**(B).**

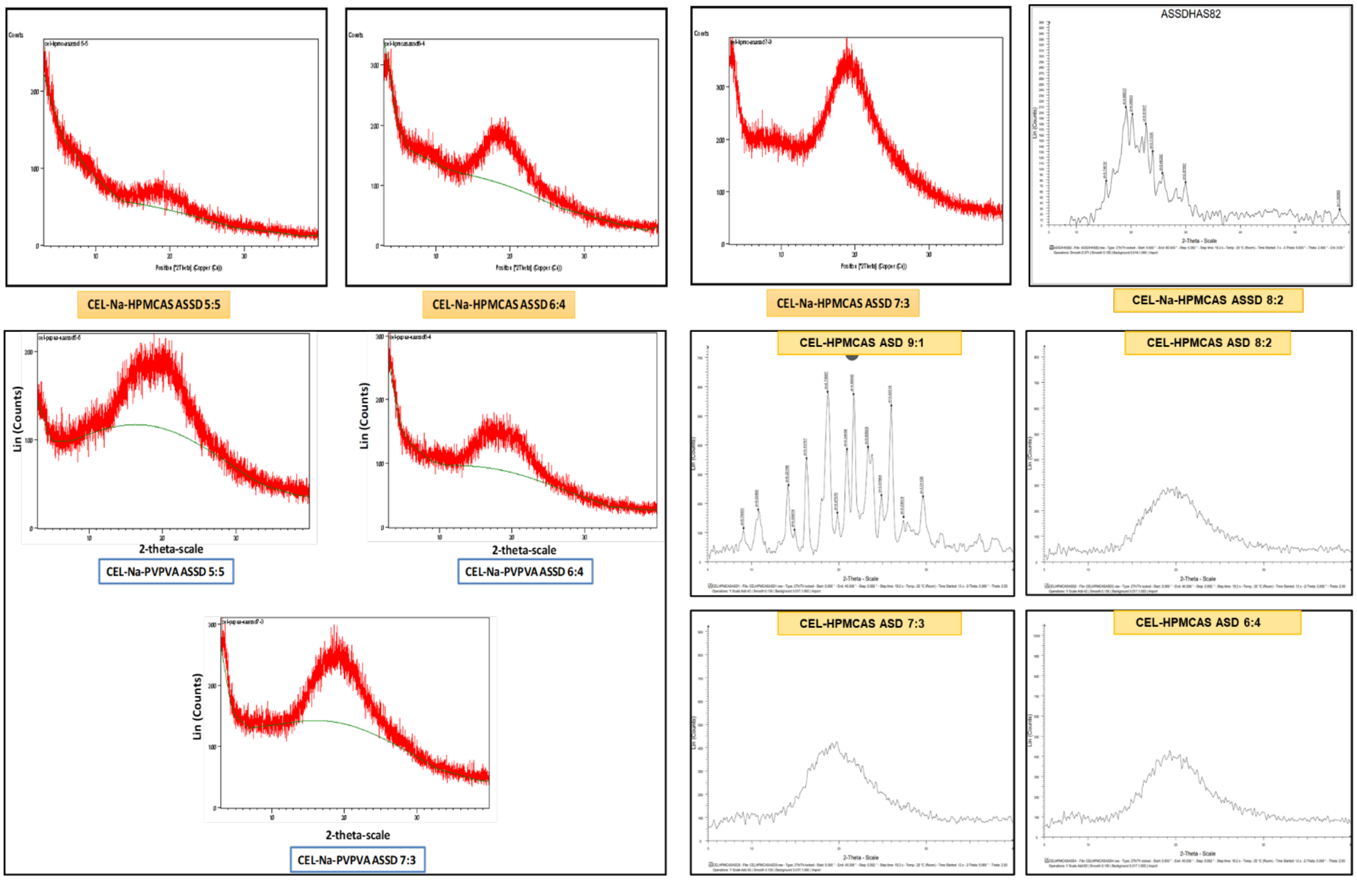


**Fig. S5.**

A. Powder X-ray diffractograms (PXRD) of crystalline CEL - showing distinct sharp peaks at different theta angles corresponding to form III - and amorphous CEL (AMR CEL), the PVP-VA and HPMCAS polymers, and the ASD and ASSD formulations at a drug-polymer ratio of 6:4 (used for all the experiments described), all showing halo patterns indicating their amorphous nature. The corresponding data is provided in Supplementary Excel File “PXRD_FigS5A_Data”. B. PXRD of CEL-HPMCAS ASDs and CEL-Na-HPMCAS and CEL-Na-PVP-VA ASSDs at drug:polymer ratios ranging from 9:1 to 5:5. A halo pattern, indicating their amorphous nature, is observed at most ratios whereas peaks indicating a crystalline form are observed for the CEL-HPMCAS ASD with a 9:1 drug:polymer ratio and the CEL-Na-HPMCAS ASSD at an 8:2 drug:polymer ratio, indicating insufficient polymer to maintain an amorphous form.

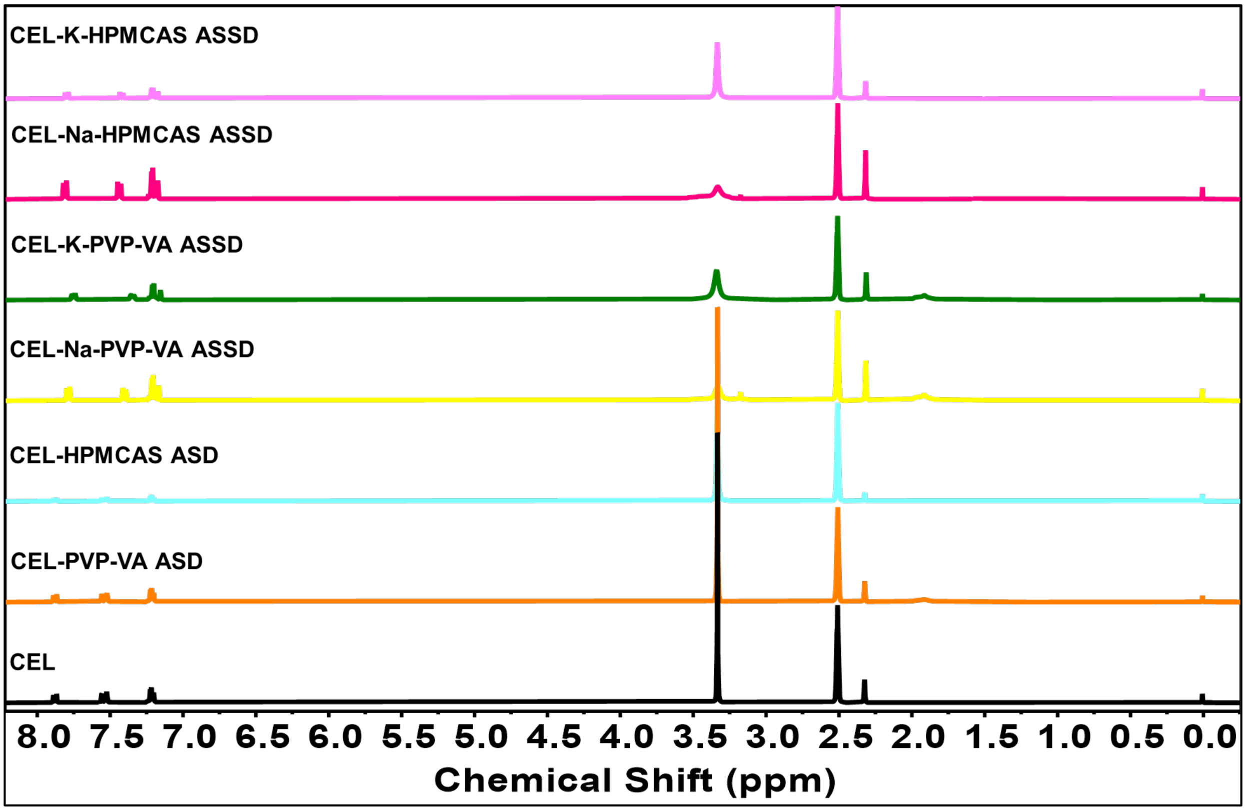


**Fig. S6**

Full NMR spectra of CEL, CEL-PVP-VA ASD, CEL-HPMCAS ASD, CEL-Na-PVP-VA ASSD, CEL-K-PVP-VA ASSD, CEL-Na-HPMCAS ASSD and CEL-K-HPMCAS ASSD showing chemical shift values on the 0 to 8.5 ppm scale. A close-up of the spectra between 7.1 and 7.9 ppm is shown in Fig. 2E.

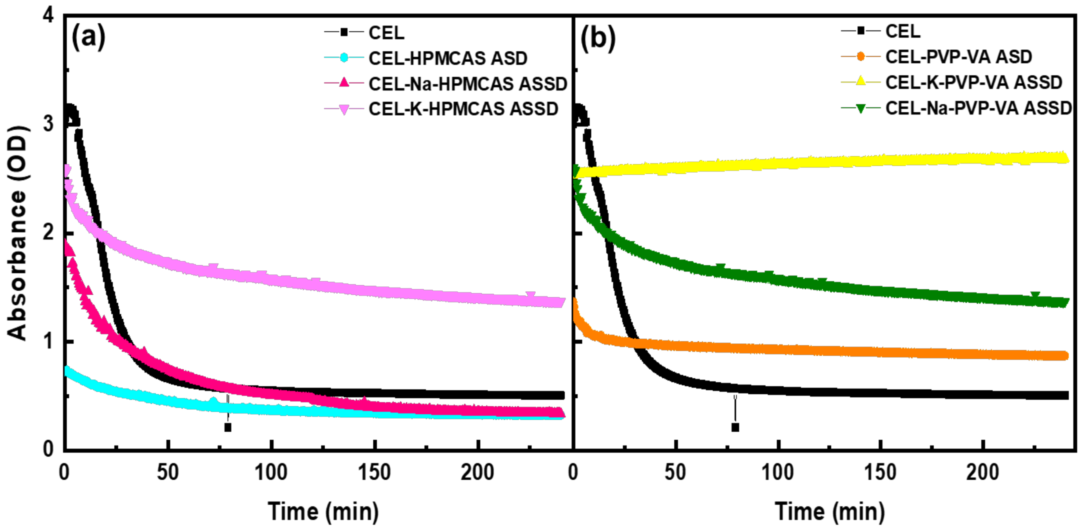


**Fig. S7**

Comparison of supersaturation and precipitation curves for crystalline CEL, and its ASD and ASSD formulations with (a) HPMCAS and (b) PVP-VA. The experiments show higher maintenance of supersaturation in ASSDs than in ASDs, with the CEL-PVP-VA ASSD having the highest ability to maintain supersaturation.

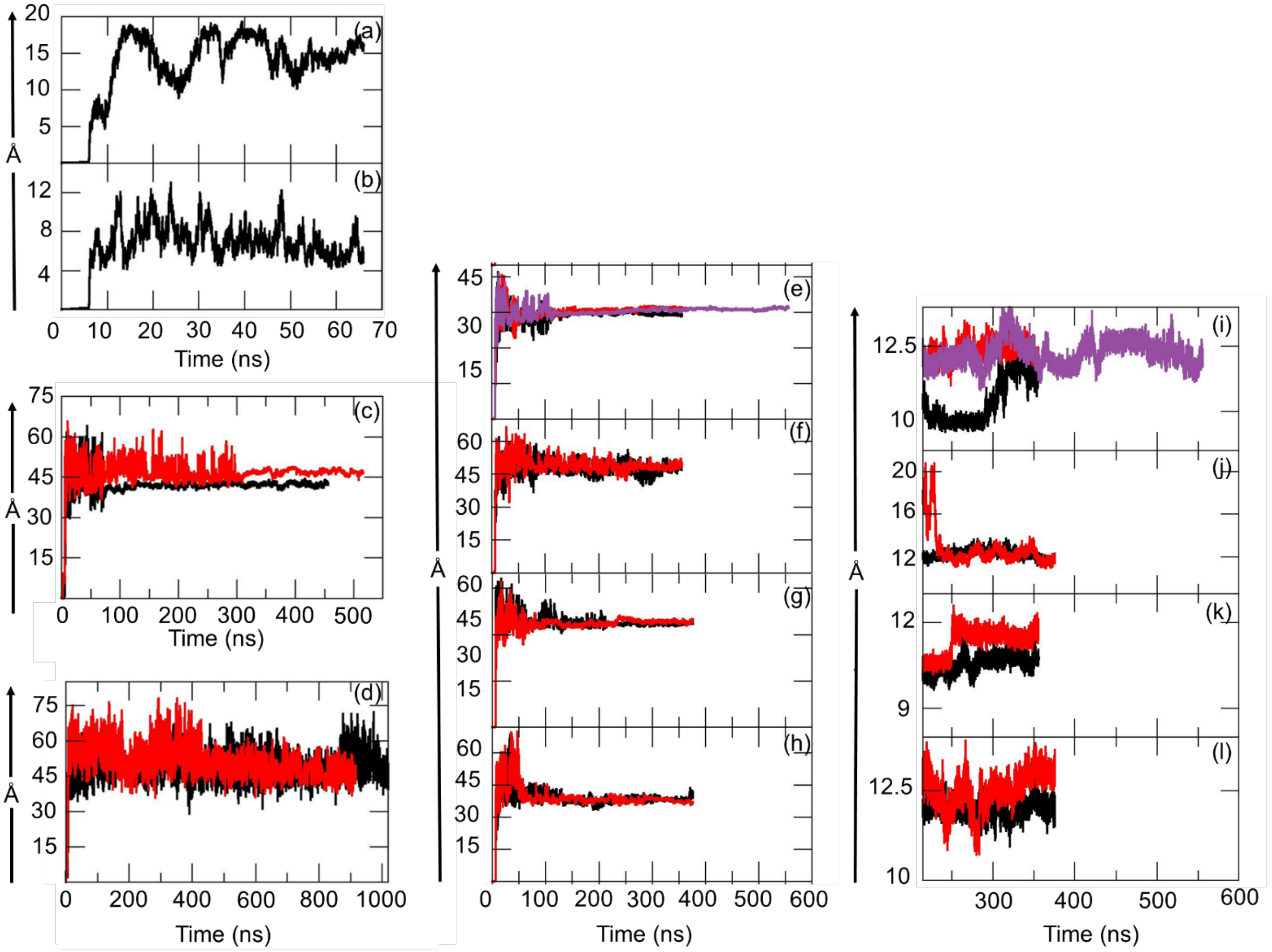


**Fig. S8**

Structural evolution of the drug and drug-polymer systems during MD simulations. All-atom root mean squared deviation (RMSD) in Å of different components of the simulated systems as a function of simulation time: (a) PVP-VA; (b) HPMCAS; (c) CEL neutral form; (d) CEL anionic form; (e) $CEL_{neutral}$-PVP-VA complex; (f) $CEL_{anion}$-PVP-VA complex; (g) $CEL_{neutral}$-HPMCAS complex; (h) $CEL_{anion}$-HPMCAS complex. Radius of gyration of the oligomer chains as a function of simulation time for the equilibrated part of the simulations of the following systems: (i) $CEL_{neutral}$-PVP-VA complex;(j) $CEL_{anion}$-PVP-VA complex;(k) $CEL_{neutral}$-HPMCAS complex; and (l) $CEL_{anion}$-HPMCAS complex. The replica simulations are colored as: replica-1 (black), replica-2 (red) and replica-3 (purple).

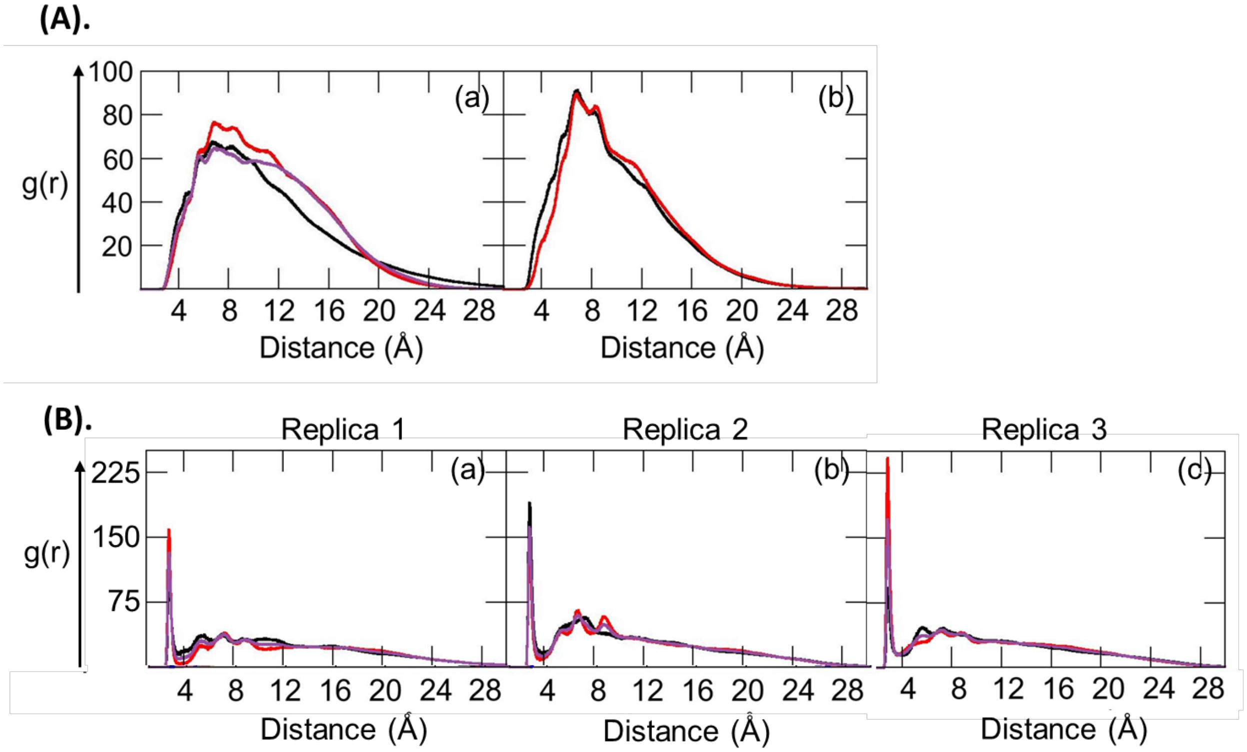


**Fig. S9**

Intermolecular CEL:PVP-VA radial distribution functions (RDFs) between computed from all-atom explicit solvent molecular dynamics simulations (replica-1: black, replica-2: red, replica-3: purple). (A) For all the fluorine groups of CEL and all the carbonyl carbon atoms of the PVP-VA oligomer for the (a) neutral form and(b) anionic form of CEL. (B) For the sulfonamide -$NH_2$ group of the neutral form of 13 CEL molecules and the carbonyl oxygen atoms of vinyl acetate (black) and vinylpyrrolidone (red) moieties and all oxygen atoms (purple) of the PVP-VA oligomer for the three replica simulations. See Fig 4 for RDFs of other systems simulated.

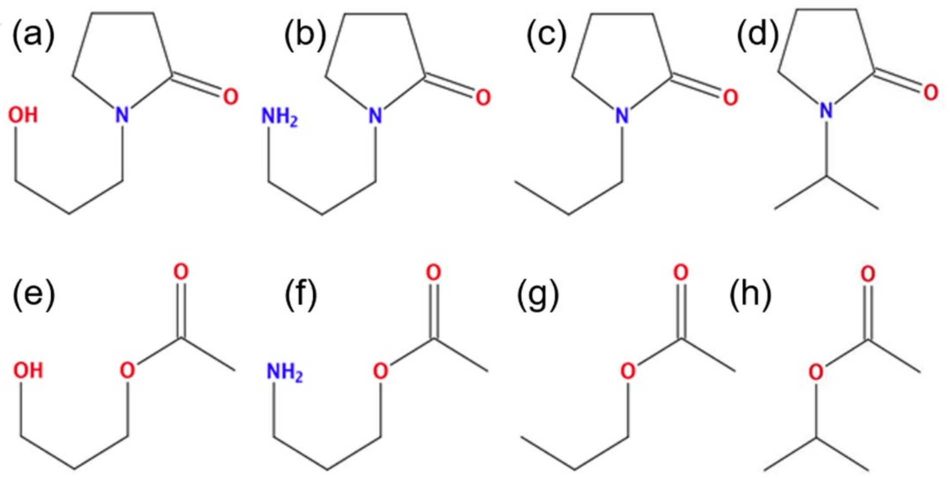


**Fig. S10**

Eight monomeric units built with the Schrödinger software (version 2020r4) to compute the electrostatic potentials (ESPs) of the non-terminal (a - c) VP and (e - g) VA repeating units and the terminal (d) VP and (h) VA units.

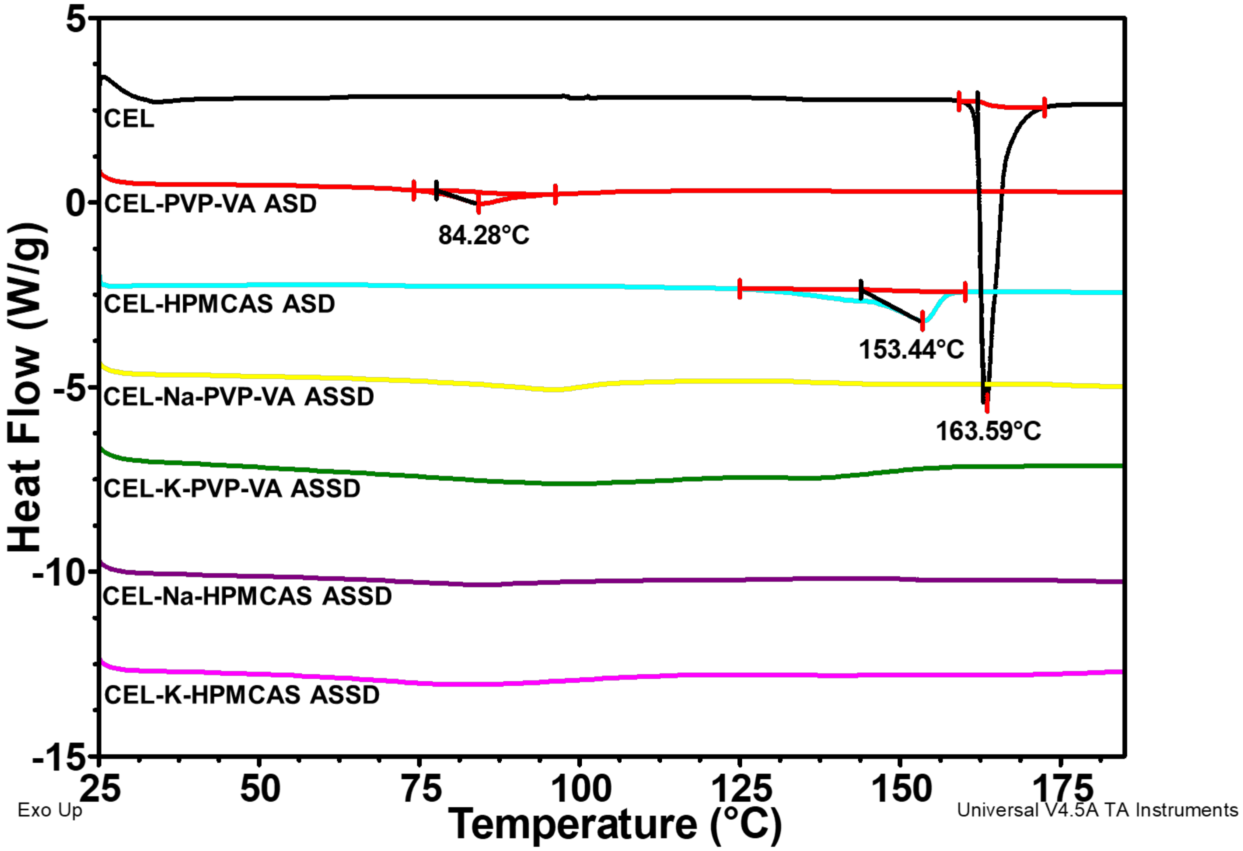


**Fig. S11**

DSC thermograms of stability samples of CEL and its amorphous formulations at ambient storage conditions after 11 months, crystalline CEL, CEL-PVP-VA ASD, CEL-HPMCAS ASD, CEL-Na-PVP-VA ASSD, CEL-K-PVP-VA ASSD, CEL-Na-HPMCAS ASSD, and CEL-K-HPMCAS ASSD. The thermograms show a single sharp melting peak for crystalline CEL, CEL-PVP-VA ASD and CEL-HPMCAS ASD at 163.59 °C, 84.28 °C and 153.44 °C, respectively, whereas no endotherm is observed in the amorphous formulations containing counterions which displayed higher $T_g$ values than their respective ASDs as shown in fig. S4. The DSC thermograms show that the amorphous nature of the ASSD but not the ASD formations was retained after 11 months, supporting the good physical stability of the ASSDs.

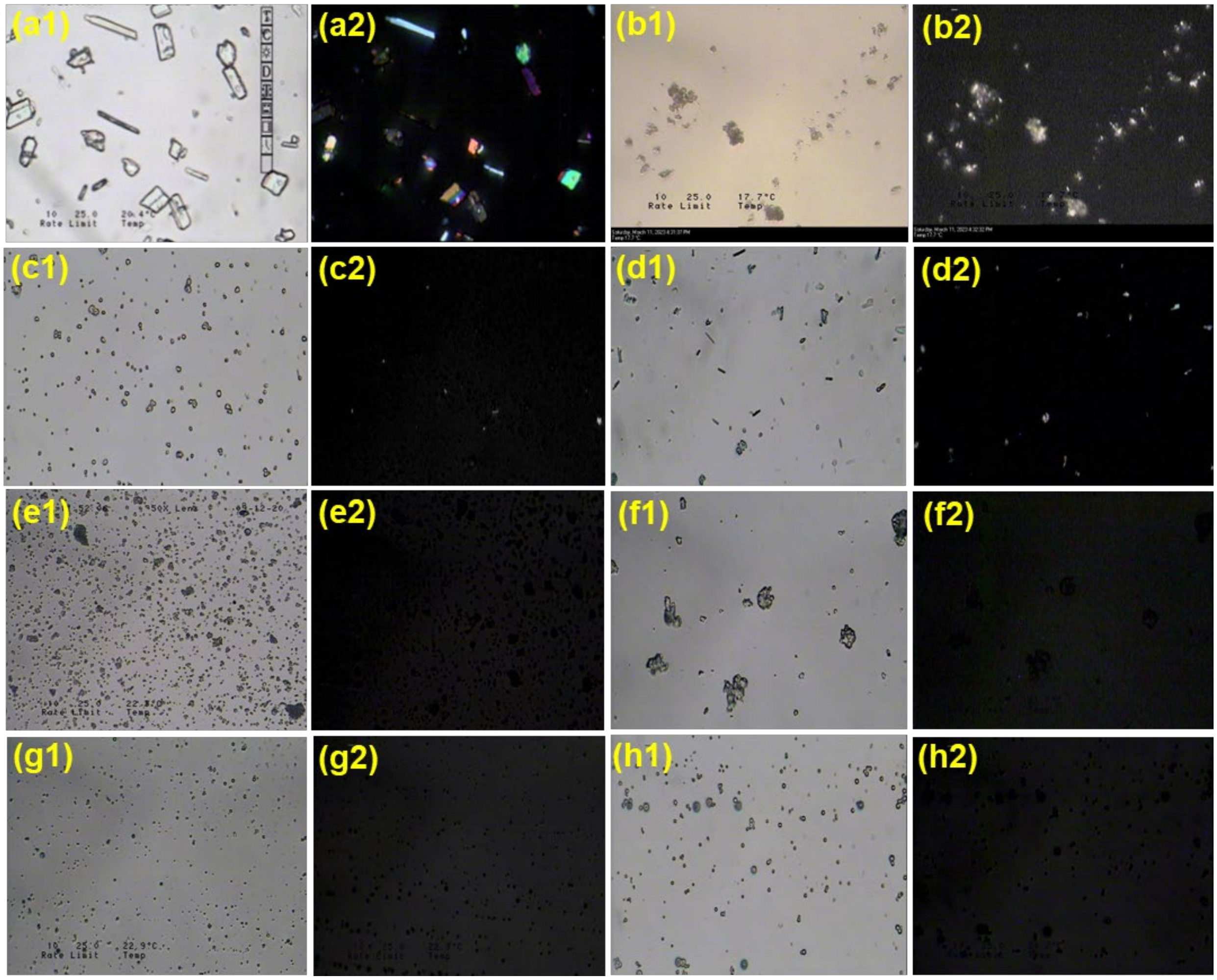


**Fig. S12**

Optical and polarised light microscopy of stability samples of CEL and its amorphous formulations at ambient storage conditions after 11 months, (a) crystalline CEL, (b) amorphous (AMR) CEL (after 48 h), (c) CEL-PVP-VA ASD, (d) CEL-HPMCAS ASD, (e) CEL-Na-PVP-VA ASSD, (f) CEL-K-PVP-VA ASSD, (g) CEL-Na-HPMCAS ASSD, and (h) CEL-K-HPMCAS ASSD (where 1 is optical mode and 2 is polarised mode). The polarised mode showed birefringence for crystalline CEL, CEL-PVP-VA ASD and CEL-HPMCAS ASD after 11 months and AMR CEL after 48 h, indicating the loss of the amorphous nature and occurrence of recrystallisation, and thus the poor stability of ASDs at longer periods. No birefringence was observed in the amorphous formulations containing counterions (e-h), indicating that the amorphous nature was retained, and recrystallization inhibited in the ASSD formulations after 11 months, thus supporting the good physical stability of ASSDs over prolonged times.

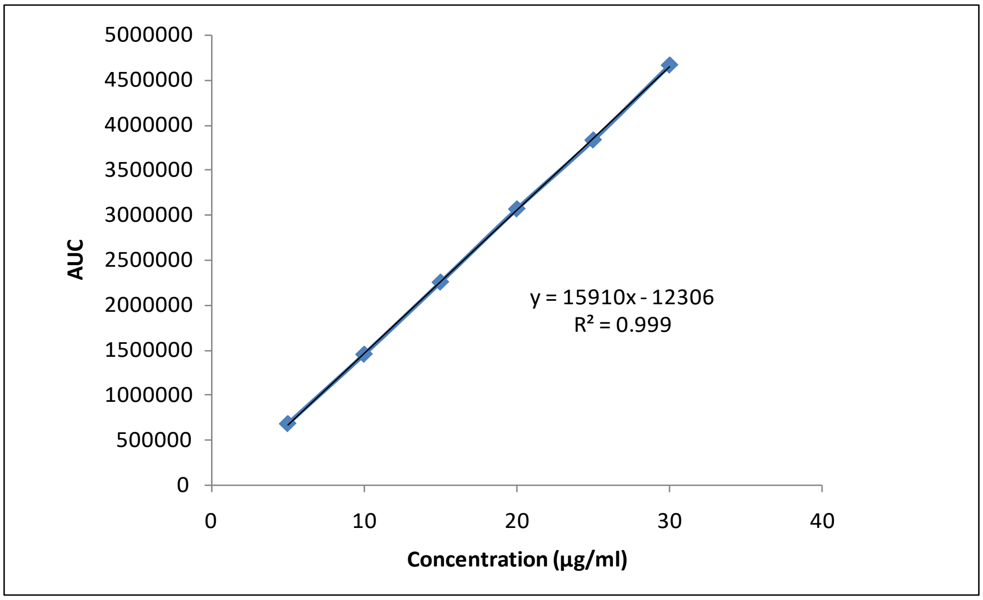


**Fig. S13.**
Validation of analytical HPLC method: Calibration plot showing linearity over a range of CEL concentrations from 5-30 µg/ml.
First, specificity, the ability to assess unequivocally the analyte in the presence of the components which may be expected to be present (impurities, degradants and matrix), was assessed. The method was found to be specific in the presence of the excipient to be used in the ASSD and no interference was observed. Next, linearity, the ability to obtain test results that are directly proportional to the concentration of analyte in the sample, was assessed. Calibration curves for the analytical method (for *in vitro* studies) were constructed by linear regression by plotting peak area versus concentration. The calibration curves were accepted only if the coefficient of correlation ($R^2$) was equal to or greater than 0.98. The linearity of the method was checked by analyzing standard solutions (at least 6 concentrations) of CEL prepared with methanol from the standard stock solution, in the working range of 5 to 100 µg/mL. The method was found to be linear.

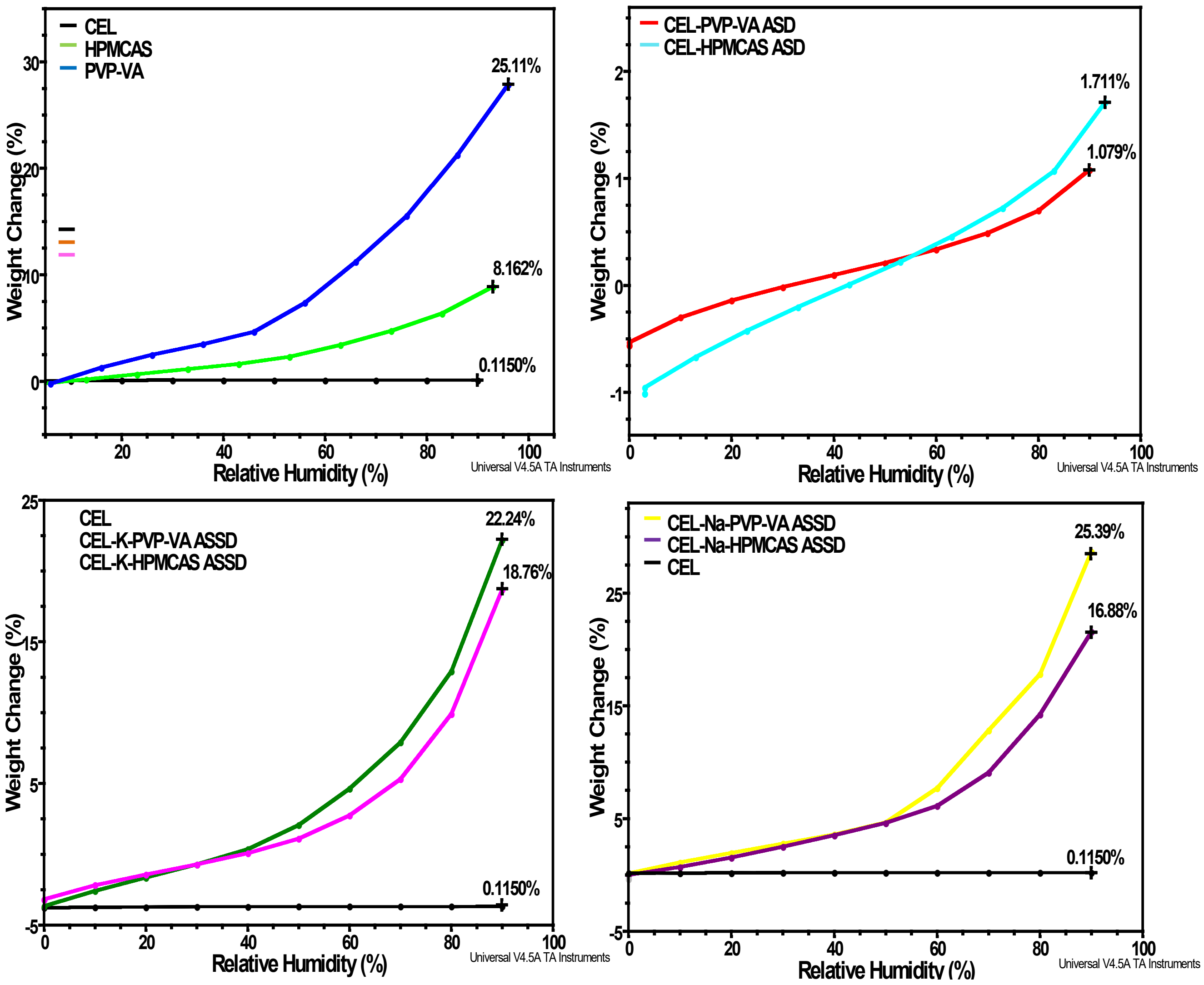


**Fig. S14.**
Moisture sorption isotherms of crystalline CEL, the PVP-VA and HPMCAS polymers, and all the amorphous ASD and ASSD formulations. The plots show more moisture sorption by the ASSD formulations than crystalline CEL and the binary ASDs at 90 % relative humidity (RH). This difference is attributed to the higher hygroscopicity of the counterions present in the ASSD formulations.

**Table S1. Computed MM/GBSA energies for CEL:CEL (CEL-CEL) and CEL:oligomer (CEL-POL) interactions in aggregates formed during MD simulations.** All values are given in kcal/mol. Values in parentheses and square brackets are for second and third replica simulations, respectively.

| CEL form | Polymer | Trajectory (ns) [a] | $\Delta G^{total}_{CEL:CEL}$ [f] | $\Delta E^{el}_{CEL:CEL}$ | $\Delta E^{vdW}_{CEL:CEL}$ | $\Delta G^{solv}_{CEL:CEL}$ | $\Delta G^{total}_{CEL:POL}$ [f] |
|---|---|---|---|---|---|---|---|
| Neutral | None | 250 - 456<br>(380 - 516) | -2.22±0.62<br>(-1.95±0.54) | -0.43±0.36<br>(-0.40±0.31) | -2.88±0.72<br>(-2.52±0.61) | 1.09±0.37<br>(0.96±0.31) | --- |
| | PVP-VA[b] | 215 – 356<br>(215 – 356)<br>[215 – 356]<br>[215 – 556] | -1.61±0.69<br>(-1.51±0.48)<br>[-1.70±0.62]<br>[-1.71±0.60] | -0.31±0.30<br>(-0.25±0.24)<br>[-0.31±0.28]<br>[-0.33±0.29] | -2.09±0.84<br>(-1.95±0.57)<br>[-2.20±0.72]<br>[-2.22±0.70] | 0.79±0.37<br>(0.69±0.27)<br>[0.81±0.32]<br>[0.84±0.33] | -8.27±9.25<br>(-11.91±7.06)<br>[-9.72±6.67]<br>[-9.91±7.07] |
| | HPMCAS | 215 – 376<br>(215 – 376) | -1.73±0.70<br>(-1.80±0.73) | -0.27±0.31<br>(-0.36±0.34) | -2.24±0.85<br>(-2.32±0.77) | 0.79±0.39<br>(0.89±0.0.40) | -9.96±6.48<br>(-9.19±7.52) |
| Anionic | None[c] | 700 - 916 | -0.95±0.47 | 15.97±3.46 | -1.73±0.70 | -15.19±3.20 | --- |
| | PVP-VA[d] | 215 – 356<br>(215 – 356) | -0.63±0.44<br>(-0.66±0.34) | 15.37±3.64<br>(16.26±3.14) | -1.24±0.65<br>(-1.27±0.51) | -14.76±3.44<br>(-15.64±2.96) | -12.14±7.75<br>(-13.62±6.99) |
| | HPMCAS[e] | 215 – 376<br>(215 – 376) | -1.04±0.56<br>(-0.91±0.50) | 20.54±3.13<br>(19.10±3.30) | -2.06±0.87<br>(-1.73±0.73) | -19.52±2.75<br>(-18.28±3.05) | -6.13±7.26<br>(-6.73±4.94) |

[a] Interaction energies were computed for frames extracted at 100 ps intervals in the time range given after the system had equilibrated. The values given are the average over all these frames and all CEL molecules in aggregates for each trajectory. Average total interaction energies are plotted in Fig. 5.

[b]For the $CEL_{neutral}$-PVP-VA system, a third replica was simulated and MMGBSA calculations were done for the two time intervals indicated. The energy values obtained for the time to 356 ns are very similar to those for the time to 556 ns, consistent with the converged behavior shown in Fig. S8 and the stability of the complex of the oligomer and the CEL molecules formed.

[c]For estimating CEL-CEL energies of anionic forms of CEL, we considered only 9 CEL molecules since, the remaining 4 CEL molecules were not part of the aggregate formed in the trajectory for system e. No CEL-CEL energies were computed for system f as the CEL molecules did not aggregate during the trajectory.

[d]For estimating CEL-CEL and CEL-PVP-VA energies, 11 anionic CEL molecules were considered and the remaining two CEL molecules that were not part of the aggregate formed were neglected in both replica trajectories.

[e]For estimating CEL-CEL and CEL-HPMCAS energies, 10 anionic CEL molecules were considered and the remaining single CEL molecule that was not part of the aggregate in the replica-1 trajectory was neglected, whereas, in the replica-2 trajectory, all the CEL molecules were considered.

[f]P-values were computed (using a 2-tail two-sample unequal variance T-test) for the differences in interaction free energies as shown in Fig. 5. The computed p-values are as follows with the values for neutral CEL given first and anionic CEL second: CEL:CEL in the absence of polymer vs. in the presence of PVP-VA – 0.02/0.04 - or HPMCAS: 0.02/0.03; CEL:PVP-VA vs. CEL:HPMCAS – 0.71/0.04.

**Table S2. MM/GBSA energy component analysis for the CEL:PVP-VA oligomer and CEL:HPMCAS oligomer complexes.** All values are given in kcal/mol. Values in parentheses and square brackets are for second and third replica simulations, respectively.

| System | Trajectory (ns) [a] | $\Delta E^{el}$ | $\Delta E^{vdW}$ | $\Delta G^{solv}$ |
|---|---|---|---|---|
| **$CEL_{Neutral}$:PVP-VA[b]** | 215 – 356<br>(215 – 356)<br>[215 – 356]<br>[215 – 556] | -2.09±3.97<br>(-2.71±3.07)<br>[-2.19±3.48]<br>[-2.24±3.58] | -9.00±9.95<br>(-12.78±7.26)<br>[-10.54±6.96]<br>[-10.65±7.34] | 2.83±3.84<br>(3.57±2.54)<br>[3.00±3.18]<br>[2.98±3.39] |
| **$CEL_{Anion}$:PVP-VA[c]** | 215 – 356<br>(215 – 356) | -9.62±7.36<br>(-10.04±6.47) | -14.31±9.22<br>(-15.78±7.83) | 11.79±7.78<br>(12.21±7.03) |
| **$CEL_{Neutral}$:HPMCAS** | 215 – 376<br>(215 – 376) | -2.91±4.81<br>(-3.10±4.91) | -13.62±7.86<br>(-12.47±9.03) | 6.58±5.30<br>(6.38±5.40) |
| **$CEL_{Anion}$:HPMCAS[d]** | 215 – 376<br>(215 – 376) | 63.29±11.53<br>(67.83±15.14) | -11.50±10.43<br>(-13.13±6.84) | -57.92±9.01<br>(-61.44±13.94) |

[a] Interaction energies were computed for frames extracted at 100 ps intervals in the time range given after the system had equilibrated. The values given are the average over all these frames and all CEL molecules in aggregates for each trajectory.
[b]For the $CEL_{neutral}$-PVP-VA system, a third replica was simulated and MMGBSA calculations were done for the two time intervals indicated. The energy values obtained for the time to 356 ns are very similar to those for the time to 556 ns, consistent with the converged behavior shown in Fig. S8 and the stability of the complex of the oligomer and the CEL molecules formed.
[c]For estimating CEL-PVP-VA energies, 11 anionic CEL molecules were considered and the remaining two CEL molecules that were not part of the aggregate formed were neglected in both replica trajectories.
[d]For estimating CEL-HPMCAS energies, 10 anionic CEL molecules were considered and the remaining single CEL molecule that was not part of the aggregate in the replica-1 trajectory was neglected, whereas, in the replica-2 trajectory, all the CEL molecules were considered.

**Table S3. Summary of 10000 bootstrapped experiments against drug:drug and drug:oligomer total binding free energies ($\Delta G^{total}$) computed by the MMGBSA method for frames from the MD simulations.** For comparison, mean, median, standard deviation (S.D.) and standard errors of the mean (SEM) values (in kcal/mol) for the full sets of frames are also given in parentheses. The corresponding p-values are < 0.05 in all but five simulations for the majority of the 10000 bootstrapped sampled data. Although for the remaining 5 simulations, the p-value was >0.05, the bootstrapped mean, median, standard deviation, and SEM were almost identical to the values for the full sets of frames, suggesting that there is no strong evidence of a difference between the groups.

| System | Mean | Median | S.D. | SEM | # of *p-values at <0.05 * |
|---|---|---|---|---|---|
| **CEL only** | | | | | |
| R1: $CEL_{Neutral}$:$CEL_{Neutral}$ | -1.95 (-1.95) | -1.92 (-1.92) | 0.54 (0.54) | 0.01 (0.01) | 9927 |
| R2: $CEL_{Neutral}$:$CEL_{Neutral}$ | -2.22 (-2.22) | -2.18 (-2.18) | 0.62 (0.62) | 0.01 (0.01) | 9999 |
| R1: $CEL_{Anion}$:$CEL_{Anion}$ | -0.95 (-0.95) | -0.97 (-0.96) | 0.47 (0.47) | 0.01 (0.01) | 6803 |
| **HPMCAS** | | | | | |
| R1: $CEL_{Neutral}$:$CEL_{Neutral}$ | -1.73 (-1.73) | -1.69 (-1.69) | 0.70 (0.70) | 0.01 (0.01) | 9936 |
| R2: $CEL_{Neutral}$:$CEL_{Neutral}$ | -1.80 (-1.80) | -1.74 (-1.74) | 0.73 (0.73) | 0.01 (0.01) | 9997 |
| R1: $CEL_{Neutral}$: HPMCAS | -9.96 (-9.96) | -10.11 (-10.10) | 6.48 (6.48) | 0.08 (0.08) | 4131 |
| R2: $CEL_{Neutral}$: HPMCAS | -9.19 (-9.19) | -7.95 (-8.00) | 7.52 (7.52) | 0.09 (0.09) | 10000 |
| R1: $CEL_{Anion}$:$CEL_{Anion}$ | -1.04 (-1.04) | -1.04 (-1.04) | 0.56 (0.56) | 0.01 (0.01) | 1741 |
| R2: $CEL_{Anion}$:$CEL_{Anion}$ | -0.91 (-0.91) | -0.90 (-0.90) | 0.50 (0.50) | 0.01 (0.01) | 7531 |
| R1: $CEL_{Anion}$: HPMCAS | -6.13 (-6.13) | -3.94 (-3.90) | 7.26 (7.26) | 0.09 (0.09) | 10000 |
| R2: $CEL_{Anion}$: HPMCAS | -6.73 (-6.73) | -6.46 (-6.50) | 4.93 (4.94) | 0.06 (0.06) | 9963 |
| **PVPVA** | | | | | |
| R1: $CEL_{Neutral}$:$CEL_{Neutral}$ | -1.61 (-1.61) | -1.63 (-1.63) | 0.69 (0.69) | 0.01 (0.01) | 7824 |
| R2: $CEL_{Neutral}$:$CEL_{Neutral}$ | -1.51 (-1.51) | -1.50 (-1.50) | 0.48 (0.48) | 0.01 (0.01) | 1875 |
| R3: $CEL_{Neutral}$:$CEL_{Neutral}$ | -1.71 (-1.71) | -1.65 (-1.65) | 0.60 (0.60) | 0.004 (0.004) | 10000 |
| R1: $CEL_{Neutral}$:PVPVA | -8.27 (-8.27) | -3.97 (-4.00) | 9.25 (9.25) | 0.11 (0.11) | 10000 |

| | | | | | |
|---|---|---|---|---|---|
| R2: $CEL_{Neutral}$:PVPVA | -11.91 (-11.91) | -10.8 (-10.8) | 7.06 (7.06) | 0.08 (0.08) | 10000 |
| R3: $CEL_{Neutral}$:PVPVA | -9.91 (-9.91) | -8.80 (-8.80) | 7.07 (7.07) | 0.05 (0.05) | 10000 |
| R1: $CEL_{Anion}$:$CEL_{Anion}$ | -0.63 (-0.63) | -0.62 (-0.62) | 0.44 (0.44) | 0.01 (0.01) | 4940 |
| R2: $CEL_{Anion}$:$CEL_{Anion}$ | -0.66 (-0.66) | -0.71 (-0.71) | 0.34 (0.34) | 0.00 (0.00) | 10000 |
| R1: $CEL_{Anion}$: PVPVA | -12.14 (-12.14) | -12.1 (-12.1) | 7.75 (7.75) | 0.10 (0.10) | 686 |
| R2: $CEL_{Anion}$:PVPVA | -13.61 (-13.62) | -14.09 (-14.10) | 6.99 (6.99) | 0.09 (0.09) | 10000 |

*P-values were computed against the median of the binding free energy values

**Table S4**. **High performance liquid chromatography (HPLC) method development**: Parameters for HPLC method for analytical estimation of CEL

| Parameters | Specifications |
|---|---|
| Mobile phase | Acetonitrile:Phosphate buffer (pH 3); 65:35 v/v |
| Type of method | Isocratic elution |
| Flow rate | 1 mL/min |
| Injection volume | 20 µl |
| Column temperature | 25°C |
| Run time | 10 min |
| Detector | PDA (UV-VIS) |
| Analytical column | Agilent TC-C18 (particle size: 5 µm, 250 mm × 4.6 mm, |

The concentration of CEL in all the samples from solubility and dissolution studies were analyzed using a validated HPLC method. The HPLC system (Shimadzu Corporation, Kyoto, Japan) included a controller (CBM-20A), a degasser (DGU-20A5), a pump (LC-20AT), an autosampler (SIL-20AC HT), a column oven (CTO-10AS VP), and a PDA detector (SPDM10AVP) with LC solution software. The reversed phase C18 column Agilent TC-C18 (250 X 4.6 mm, 5 µm, Agilent technologies) maintained at 25 °C was used for HPLC analysis. The mobile phase, consisting of acetonitrile and phosphate buffer (10 mM, pH 3) in a ratio of 65:35 v/v, was used and pumped isocratically at 1 mL/min flow rate. A wavelength of 252 nm was set in the PDA detector.

For analysis of samples from *in vivo* pharmacokinetic studies, the bioanalytical method was developed with minor modifications. All the samples were analyzed with the above mobile phase in a ratio of 55:45 v/v. Indomethacin was used as an internal standard. The injection volume was 40 µL. A primary stock solution of CEL (100 µg/mL) was prepared by dissolving 1 mg of CEL in 10 mL of HPLC grade methanol and further diluted to the desired concentration ranges from 5 to 100 µg/mL for preparing standard calibration curve. Phosphate buffer (10 mM, pH 3) was prepared by dissolving 870 mg of di-potassium hydrogen orthophosphate in 500 mL of distilled water and adjusting the pH with orthophosphoric acid. This buffer was filtered through a 0.45 µm membrane filter (Nylon) and degassed ultrasonically.

**Table S5. Precision and accuracy of the HPLC method**. Intra- and inter-day accuracy and precision data for three different concentrations of CEL.

| Concentration (µg/ml) | Accuracy | | Precision | |
|---|---|---|---|---|
| | Intraday | Interday | Intraday | Interday |
| 16 | 100.36±1.8 | 100.41±1.35 | 1.54±0.32 | 0.89±0.66 |
| 20 | 99.73±2.36 | 98.12±2.28 | 1.83±0.59 | 0.95±1.28 |
| 24 | 98.29±1.14 | 99.68±0.87 | 1.16±0.59 | 0.97±0.65 |

The precision of the analytical method expresses the closeness of agreement between a series of measurements obtained from multiple sampling of the same homogeneous sample. The accuracy of the analytical method represents the closeness of test results obtained by the method to the nominal (true) value of analyte. Intra-day accuracy and precision were evaluated by analysis of quality control (QC) samples at different times on the same day. Inter-day accuracy and precision were calculated from results of replicate assays of QC samples. The concentration of each sample was determined by use of calibration standards prepared on the same day. The accuracy of the method was estimated by measurement of recovery, and precision was estimated by calculating the relative standard deviation, RSD. Accuracy was assessed using a minimum of nine (n=9) determinations over a minimum of three (n=3) concentration levels covering the specified range (3 concentrations/3 replicates), whereas precision was analyzed in terms of repeatability and obtained by calculating the relative standard deviation of nine determinations over a minimum of 3 concentration levels covering the specified range. Both parameters were found to be in the specified range as per [Q2 (R1)] ICH guidelines.

The limit of detection (LOD) was determined as the lowest concentration in the standard calibration curve which gave a signal-to-noise ratio higher than 3. The limit of quantification (LOQ) was determined as the concentration which gave a signal-to-noise ratio higher than 5. LOQ and LOD were determined from the slope and the standard deviation of the y-intercept of the standard plot equation from formulas F1 and F2, and were found to be 0.35 and 1.0 µg/mL, respectively.

$$\boldsymbol{LOD} = \frac{\boldsymbol{SD}}{\boldsymbol{Slope}} \times \mathbf{3.3} \qquad \ldots\ldots\ldots\ldots\ldots\ldots\ldots\ldots\ldots\ldots\ldots\ldots\ldots\ldots\ldots (\boldsymbol{F1})$$

$$\boldsymbol{LOQ} = \frac{\boldsymbol{SD}}{\boldsymbol{Slope}} \times \mathbf{10} \qquad \ldots\ldots\ldots\ldots\ldots\ldots\ldots\ldots\ldots\ldots\ldots\ldots\ldots\ldots\ldots\ldots (\boldsymbol{F2})$$
